# Extracellular Matrix-Induced Genes May Reduce Response to Rapamycin in LAM

**DOI:** 10.1101/2024.05.16.594484

**Authors:** D Clements, R Babaei-Jadidi, J Johnson, S Miller, N Shah, JMB Sand, DJ Leeming, LA Borthwick, AJ Fisher, A Dufour, SR Johnson

## Abstract

**Rationale:** Lymphangioleiomyomatosis (LAM) is a rare cystic lung disease driven by nodules containing TSC2^-/-^ ‘LAM cells’ and recruited LAM associated fibroblasts (LAFs). Although rapamycin reduces lung function loss, some patients continue to decline meaning additional therapies are needed.

**Objectives:** To investigate how the LAM nodule environment affects LAM cell proliferation and the response to rapamycin.

**Methods:** Changes in advanced LAM were identified using shotgun proteomics and immunohistochemistry in tissue from carefully phenotyped patients. Genes potentially associated with rapamycin insensitivity of cells grown on LAF-derived extracellular matrix were identified by RNA sequencing and validated using repurposed pharmacologic inhibitors.

**Main Results:** More advanced disease was associated with increasing nodules adjacent to lung cysts and greater decline in forced expiratory volume in 1 sec (FEV_1_) when treated with rapamycin (p=0.005). In late-stage LAM, proteomics identified upregulation of pathways associated with accumulation of activated fibroblasts, including extracellular matrix deposition, glucose metabolism and the actin cytoskeleton. Picrosirius red staining and immunohistochemistry confirmed deposition of extracellular matrix within LAM nodules. The growth of TSC2^-/-^ model LAM cells was increased on LAF-derived extracellular matrix (LAF ECM), and incompletely supressed by rapamycin (p<0.0001). RNA sequencing of cells grown on LAF ECM identified upregulation of pathways driving cell cycle control, transcription and metabolism in cells. Tractable, pro-proliferative, rapamycin insensitive genes included *CDK7*, *GAS6* and *PLAU.* Repurposed inhibitors of these pathways inhibited LAM cell proliferation and enhanced the anti-proliferative effect of rapamycin.

**Conclusions:** Extracellular matrix deposited by LAM associated fibroblasts upregulates expression of genes which potentially blunt the response to rapamycin, but offer additional therapeutic opportunities for patients with established LAM.

## Introduction

Lymphangioleiomyomatosis (LAM) is a rare, almost entirely female-specific multisystem disease [1]. The pulmonary features of LAM include progressive cystic changes to the lung parenchyma associated with aggregates or nodules of cells; as a result, patients suffer an accelerated loss of lung function which can result in respiratory failure. In the lung, pathogenic changes are initiated by a cell of unknown origin termed a ‘LAM cell’, which has undergone biallelic loss of *TSC2* gene function, or more rarely loss of *TSC1* [2]. These genes encode the tumour suppressor proteins tuberin and hamartin, which together regulate the activity of the mTOR kinase-containing complex 1, MTORC1. In the absence of tuberin or hamartin function MTORC1 is aberrantly activated, potentially resulting in increased proliferation and metabolic changes in LAM cells. LAM cells in the parenchyma cause degradation of the alveolar structure by a mechanism as yet unknown, but hypothesised to be a consequence of dysregulated extracellular matrix (ECM)-degrading protease expression [3,4]. There is also recruitment of other cell types, including fibroblasts and mast cells [5,6], resulting in the formation LAM nodules. The contribution of these stromal cells to disease progression is unclear.

The identification of mTORC1 as the cellular target of tuberin and hamartin led to the repurposing of the immunosuppressive and antifungal drug rapamycin as a treatment for LAM [7]. For many patients, rapamycin is an effective and tolerable treatment, stabilising lung function as long as treatment is continued [8]. Cessation of treatment results in continued disease progression, and rapamycin is considered cytostatic rather than cytotoxic. Some patients continue to lose lung function despite rapamycin treatment, and Johnson *et al*. observed a continued decline in carbon monoxide diffusing capacity (DL_CO_) in patients treated with rapamycin, even when their Forced Expiratory Volume in 1 second (FEV_1_) was stable [9]. It is not clear why some patients respond less well to rapamycin, although, in a study by Bee *et al*. [10], patients with the greatest loss of lung function while undergoing rapamycin treatment also tended to be those with longer disease duration and lower lung function at the start of the treatment, suggesting that more advanced disease may be less sensitive to rapamycin. Thus, despite the undoubted benefit that rapamycin confers to patients with LAM, additional treatments would be valuable both for patients with poor or partial response to rapamycin and to effect a cure for LAM by triggering LAM cell death.

Previously we identified fibroblasts, which we termed LAM Associated Fibroblasts, or LAFs, as a component of LAM nodules, and proposed that these cells could contribute to disease progression by supporting LAM cell growth and survival [5]. Using immunohistochemical markers of cell types, it was found that the ratio of LAFs relative to LAM cells increases in LAM nodules over time. [11]. We hypothesise that, in part, LAFs contribute to disease progression in LAM by modification of the microenvironment of LAM cells via accumulated deposition of ECM. We propose that LAF-deposited ECM supports LAM cell proliferation and survival in the presence of rapamycin, and that targeting the response of LAM cells to ECM will provide new therapeutic options for LAM patients.

In this study, proteomic analysis of late-stage LAM lung tissue showed enrichment of proteins involved in pathways associated with fibroblast activation and ECM deposition, and collagen deposition was observed in LAM lung tissue sections. *In vitro*, decellularised LAF-deposited ECM supported increased proliferation of TSC2^-/-^ LAM-derived cells, and a transcriptional analysis of these cells revealed a number of upregulated pro-proliferative pathways. Using inhibitors of these pathways we were able to effect a greater inhibition of TSC2^-/-^ cell growth *in vitro* than treatment with rapamycin alone, supporting the hypothesis that targeting the interaction between LAM cells and their microenvironment may provide additional benefit for patients.

## Methods

### Patient information and sample collection

Tissue for LAM immunohistochemistry and collagen staining was obtained from 32 patients, including explanted diseased lungs from six patients with LAM undergoing lung transplantation. This cohort was described previously (1). LAM lung tissue for proteomic analysis was obtained from women undergoing lung transplantation at Harefield Hospital, Uxbridge, UK or Freeman Hospital, Newcastle upon Tyne, UK, and was stored at -80°C until use. Use of tissue was approved by Research Ethics Committee (East Midlands REC, 13/EM/0264; North East-Newcastle and North Tyneside 1, 11/NE/0291) and all subjects provided written informed consent.

Non-transplanted healthy lungs were obtained using a tissue retrieval service (International Institute for the Advancement of Medicine, Edison, NJ) and were received 24–48 h post-surgical removal. Informed consent for organs to be harvested for transplant or for research was obtained from family members at each hospital. No personal identifying information was provided for any of the donors.

For serum isolation, blood was collected in serum separation tubes and processed within 2 h of collection; samples were allowed to clot at room temperature for 1 hour, centrifuged, and serum was stored in aliquots at − 80 °C until assayed.

### Cell culture

Primary human lung fibroblasts (LAFs) were derived from surgical biopsies and explanted diseased lung tissue by collagenase digestion as described previously [12]. All cells were maintained in phenol red free DMEM-F12 medium (Gibco, 21041-033) with 10% Foetal Calf Serum unless otherwise stated.

### Preparation of decellularised extracellular matrix

Fibroblast deposited, decellularised extracellular matrix was prepared as described [13] after 7 days of fibroblast culture, using freeze thaw cycling followed by ammonium hydroxide treatment. Plate were stored at -80°C until use.

### Cell proliferation and inhibitor assays

Proliferation assays were carried out using cell counts, the Bromodeoxyuridine (BrdU) Cell Proliferation Assay Kit (#6813, Cell Signaling Technology), resazurin reduction (alamarBlue™ Cell Viability Reagent, Invitrogen, DAL 1025) and Thiazolyl Blue Tetrazolium Bromide (MTT) reduction (M5655, Sigma-Aldrich). Inhibitors were: Samuraciclib hydrochloride (Insight Biotechnology, HY-103712A), Dubermatinib (Insight Biotechnology, HY-12963), IPR-803 (Cambridge Biosciences, HY-111192), and Rapamycin (InSolution Rapamycin, Sigma #553211). All were dissolved in DMSO, which was used as the vehicle control; the maximum concentration of DMSO in media was 0.001%. Experimental conditions were run in triplicate wells with at least three independent experiments performed.

### Immunohistochemistry and collagen staining

Paraffin embedded LAM and control lung tissues were dewaxed in xylene and rehydrated through an ethanol series (100%, 95%, 70%, water). If specified by the antibody supplier, antigen retrieval was carried out by heating sections to 100°C in a steamer for 15 minutes in 10 mM Sodium Citrate, 0.05% Tween 20, pH 6.0 or 10 mM Tris, 1 mM EDTA, 0.05% Tween 20, pH 9.0. Endogenous peroxidase activity was quenched by incubating sections in 3% H_2_O_2_ in water for 10 minutes at room temperature.

Primary antibodies were from Proteintech (Manchester, UK): AXL (13196-1-AP), GAS6 (13795-1-AP), uPAR (10286-1-AP), uPA/Urokinase (17968-1-AP), CDK7 (27027-1-AP), Collagen alpha-1(I) chain (14695-1-AP), Collagen alpha-1(VI) chain (17023-1-AP). Anti-melanoma antibody PNL2 (Zytomed 188-10864) was from 2BScientific (Oxford, UK), mouse monoclonal anti-Smooth Muscle Actin antibody (Clone 1A4) was from Sigma (A2547).

Secondary antibodies were Horse Radish Peroxidase conjugated goat anti-rabbit or anti-mouse (ImmPRESS Polymer Detection Kits, Peroxidase, Vector Laboratories, MP-7451 and MP-7452) or Dako REAL EnVision detection system (Dako, K5007). Detection of primary antibodies was performed using ImmPACT DAB Peroxidase substrate (Vector Laboratories, SK-4105). Sections were counterstained with Mayer’s haematoxylin, dehydrated through an ethanol series to xylene, and mounted in VectaMount Permanent Mounting Medium (Vector Laboratories, H-5000-60).

Picrosirius red (PSR) staining for fibrillar collagens was performed by Translational Research, Division of Cellular Pathology, Nottingham University Hospitals (NUH) Trust. Quantification of PSR staining was carried out using Fiji (https://fiji.sc/) [14].

Slides were scanned at 40x magnification on a Hamamatsu Nanozoomer slide scanner.

### Shotgun proteomics

For shotgun proteomics, lung parenchymal fragments from five LAM donors undergoing lung transplantation (**Supplementary Figure 1**) and six lungs from healthy cadaveric donors were used. Samples were lysed and reduced with 5 mM DTT (Gold Biotechnology, St-Louis, MO) at 37 °C for 1 h and alkylated with 15 mM IAA (GE Healthcare, Mississauga, ON) in the dark at room temperature for 30 min followed by quenching with 15 mM DTT. Proteins were subjected to trypsin (Promega, Madison, WI) digestion overnight at 37°C. The pH was next adjusted to 6.5 before the samples were isotopically labelled with a final concentration of 40 mM deuterated heavy formaldehyde (LAM patient samples) or 40 mM light formaldehyde (healthy control samples) in the presence of 40 mM sodium cyanoborohydride overnight at 37°C. Samples were then desalted using Sep-Pak C18 columns and lyophilized before submitting for LC-MS/MS analysis to the Southern Alberta Mass Spectrometry core facility, University of Calgary, Canada.

### High performance liquid chromatography (HPLC) and mass spectrometry

All liquid chromatography and mass spectrometry experiment were carried out by the Southern Alberta Mass Spectrometry core facility at the University of Calgary, Canada. Analysis was performed on an Orbitrap Fusion Lumos Tribrid mass spectrometer (Thermo Fisher Scientific, Mississauga, ON) operated with Xcalibur (version 4.0.21.10) and coupled to a Thermo Scientific Easy-nLC (nanoflow Liquid Chromatography) 1200 system. Tryptic peptides (2 µg) were loaded onto a C18 trap (75 µm × 2 cm; Acclaim PepMap 100, P/N 164946; Thermo Fisher Scientific) at a flow rate of 2 µL/min of solvent A (0.1% formic acid and 3% acetonitrile in LC-mass spectrometry grade water). Peptides were eluted using a 120 min gradient from 5 to 40% (5% to 28% in 105 min followed by an increase to 40% B in 15 min) of solvent B (0.1% formic acid in 80% LC-mass spectrometry grade acetonitrile) at a flow rate of 0.3% µL/min and separated on a C18 analytical column (75 µm × 50 cm; PepMap RSLC C18; P/N ES803; Thermo Fisher Scientific). Peptides were then electrosprayed using 2.3 kV into the ion transfer tube (300° C) of the Orbitrap Lumos operating in positive mode. The Orbitrap first performed a full mass spectrometry scan at a resolution of 120,000 FWHM to detect the precursor ion having a mass-to-charge ratio (m/z) between 375 and 1575and a +2 to +4 charge. The Orbitrap AGC (Auto Gain Control) and the maximum injection time were set at 4 × 10^5^ and 50 ms, respectively. The Orbitrap was operated using the top speed mode with a 3s cycle time for precursor selection. The most intense precursor ions presenting a peptidic isotopic profile and having an intensity threshold of at least 2 × 10^4^ were isolated using the quadrupole (isolation window of m/z 0.7) and fragmented with HCD (38% collision energy) in the ion routing Multipole. The fragment ions (MS2) were analysed in the Orbitrap at a resolution of 15 000. The AGC, the maximum injection time and the first mass were set at 1 ×10^5^, 105 ms, and 100 ms, respectively. Dynamic exclusion was enabled for 45 s to avoid the acquisition of the same precursor ion having a similar m/z (±10 ppm).

### Proteomic data and bioinformatic analysis

Spectral data were matched to peptide sequences in the human UniProt protein database using the MaxQuant software package v.1.6.0.1, peptide-spectrum match false discovery rate (FDR) of < 0.01 for the shotgun proteomics data. Search parameters included a mass tolerance of 20 p.p.m. for the parent ion, 0.05 Da for the fragment ion, carbamidomethylation of cysteine residues (+57.021464), variable N-terminal modification by acetylation (+42.010565Da), and variable methionine oxidation (+15.994915Da). For the shotgun proteomics data, cleavage site specificity was set to Trypsin/P (search for free N-terminus and only for lysines), with up to two missed cleavages allowed. The files evidence.txt and proteinGroups.txt were analysed by MSstatsTMT (v2.4.0) [15] using R software (v4.2.0) (R Core Team (2022). R: A language and environment for statistical computing. R Foundation for Statistical Computing, Vienna, Austria. URL https://www.R-project.org/) for the statistical analysis. Significant outlier cut-off values were determined after log(2) transformation by boxplot-and-whiskers analysis using the BoxPlotR tool. Database searches were limited to a maximal length of 40 residues per peptide. Peptide sequences matching reverse or contaminant entries were removed.

STRING v.12.0 (Search Tool for the Retrieval of Interacting Genes, https://string-db.org, [16]) was used to identify interactions among the proteins found to be upregulated greater than 1.3-fold in lung tissue from LAM lung tissue relative to normal lung tissue. Data were analysed at ‘high confidence’, with a false discovery rate (FDR) correction of 1%. Enriched pathways were identified using the automated pathway enrichment analysis, using the entire genome as statistical background, and data from Gene Ontology annotations were used to identify upregulated pathways. Enrichr (https://maayanlab.cloud/Enrichr/) [17–19], was also used to identify enriched pathways.

### COL6A1 neoepitope assay

Serum samples from LAM patients and healthy controls were analysed using the competitive ELISA (nordicC6M^TM^ , Nordic Bioscience, Denmark) detecting a fragment of Collagen alpha-1(VI) chain (COL6A1) generated by MMP-2 and MMP-9 digestion (C6M) as previously described [12].

### RNA sequencing: Library preparation, QC and Sequencing protocol

621-101 cells, a gift from Prof E. Henske, were derived from the angiomyolipoma of a woman with LAM and carry bi-allelic inactivation of the TSC2 gene [20]. These cells were grown on either tissue culture plastic or LAF deposited ECM from three LAF donors, for five days in DMEM-F12 medium with 1% Foetal Calf serum, in the presence and absence of 10nM rapamycin (InSolution Rapamycin, Sigma #553211). Each sample was triplicated. Cells were harvested and total RNA extracted using the Qiagen RNeasy kit, followed by DNA removal using RQ1 RNase-Free DNase (Promega) and Monarch RNA Clean Up Kit (NEB). Triplicated samples were pooled prior to sequencing.

#### Sample QC

RNA concentrations were measured using the Qubit Fluorometer and the Qubit RNA BR Assay Kit (ThermoFisher Scientific; Q10211) and RNA integrity was assessed using the Agilent TapeStation 4200 and the Agilent RNA ScreenTape Assay Kit (Agilent; 5067-5576 and 5067-5577).

#### Library Preparation

For all samples, cDNA was generated from 400ng of total RNA using the QuantSeq 3’ mRNA-Seq library prep kit for Illumina (FWD) (Lexogen; 015.96). Indexed sequencing libraries were prepared using the Lexogen i7 6nt Index Set (Lexogen; P04496). For all samples - 13 cycles of PCR.

#### Library QC

Libraries were quantified using the Qubit Fluorometer and the Qubit dsDNA HS Kit (ThermoFisher Scientific; Q32854). Library fragment-length distributions were analyzed using the Agilent TapeStation 4200 and the Agilent High Sensitivity D1000 ScreenTape Assay (Agilent; 5067-5584 and 5067-5585). Libraries were pooled in equimolar amounts and final library quantification performed using the KAPA Library Quantification Kit for Illumina (Roche; KK4824).

The library pool was sequenced on the Illumina NextSeq 500, on a NextSeq 500 High Output v2.5 75 cycle kit (Illumina; 20024906), to generate approximately 5 million 75bp single-end reads per sample. Data analysis was carried out using Lexogen QuantSeq Data Analysis (REV) software on the BlueBee® Platform.

The data discussed in this publication have been deposited in NCBI’s Gene Expression Omnibus ([21] https://www.ncbi.nlm.nih.gov/geo/), and are accessible through GEO Series accession number GSE265807.

### Statistical Analyses

Statistical analyses are described in the Figure legends and were performed using Graphpad Prism Version 10.1.1. For analyses comparing two groups, unpaired Student’s *t* test or Mann-Whitney test was used as indicated in the Figure legend. For more than two groups we used ANOVA or Kruskal-Wallace test as indicated. Values of p < 0.05 were considered significant between two groups. In graphs, p values less than 0.05 are given one asterisk, p values less than 0.01 are given two asterisks, p values less than 0.001 are given three asterisks, and p values less than 0.0001 are given four asterisks.

## Results

### Disease progression and response to rapamycin

Pulmonary LAM is characterised by parenchymal cysts associated with alpha Smooth Muscle Actin (αSMA)-expressing nodules or aggregates of cells, which include both LAM cells and LAM-associated stromal fibroblasts (LAFs) [5]. Examination of lung sections from patients with linked clinical information reveals the progression of LAM from the earliest stages (disease duration less than 12 months), in which cysts are evident but nodules are not, through late stage disease (disease duration greater than 12 months) with more pronounced αSMA-positive nodules, to end-stage disease (diseased lung from lung transplant patients) in which normal parenchymal structure is disrupted by an extensive nodular or reticular expansion of αSMA-positive cells **(Figure 1A)**. Previously, using immunohistochemical markers of LAM cells and fibroblasts, we showed that the composition of LAM nodules evolves as the disease progresses, with LAFs becoming more abundant with later stage disease [11].

**Figure 1:**
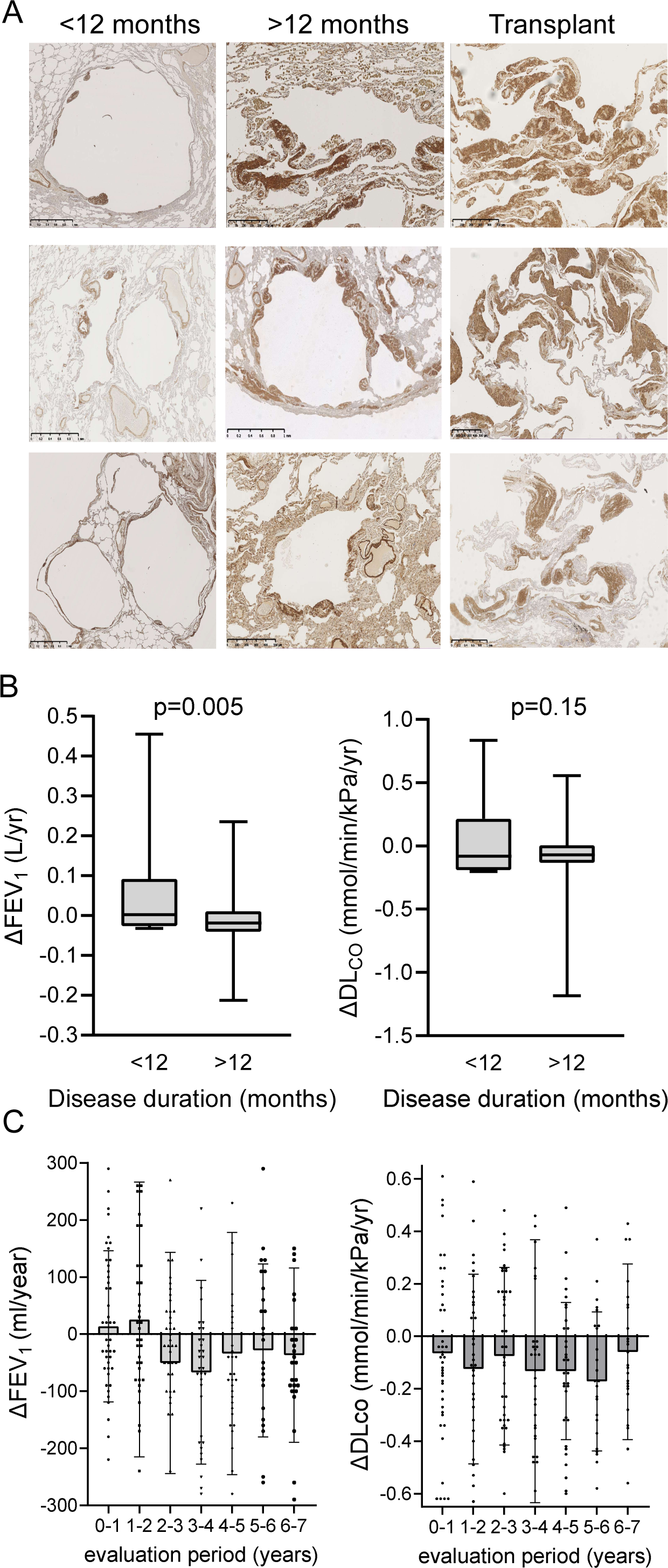
Relationship between disease duration and mTOR inhibitor response. (**A**) LAM nodules identified by immunoreactivity to anti-Smooth Muscle Actin (brown staining). Representative images from 12 patients with a first symptom of LAM <12 months and 15 patients with a disease duration of >12 months taken for clinical purposes or transplant explants. Nodules in patients with disease greater than 12 months duration are larger and more common in cyst walls than in those with very early disease. (**B**) Rate of change in lung function (Δ) in seven patients with a disease duration of <12 months and 59 patients with a disease duration of >12 months treated with rapamycin for a mean duration of analysed by unpaired t-test for ΔFEV_1_ and ΔDLco. (**C**) Annual rate of change in FEV1 and DLco for 12 months periods from the start of MTORC1 inhibitor treatment.

To determine if rapamycin response could be related to these changes we compared rate of loss of FEV_1_ and DL_CO_ in patients treated with rapamycin for active lung disease stratified by disease duration. We identified seven patients who had treatment initiated less than 12 months after symptom onset, compared with 59 patients with treatment initiated more 12 months after symptom onset. Rate of change of FEV_1_ was 7 ml/yr (95% confidence interval (CI) -8 to +23) for those treated within 12 months compared with -13 ml/yr (95% CI -3 to +2) for treatment initiated after 12 months (p=0.005) over a mean period of observation of 98 (SD 46) months. For DL_CO_, the rate of change for those treated within 12 months was 0.076 mmol/min/kPa/yr (95% CI -0.26 to +0.41) compared with -0.062 (95% CI -0.12 to -0.004) for treatment initiated after 12 months (p=0.15) (Figure 1B). Patients with shorter disease durations were a mean of 7.3 years younger (p=0.037) and had higher pre-treatment FEV_1_ values (71.3 (15.6) vs. 54.3 (20) percent predicted) but similar values for DLco (40.2 (7.8) vs. 41.7 (12.6)) compared with those with a disease duration >12 months. We next examined the annual change in lung function for each year of treatment in a national cohort of women with LAM treated with rapamycin. Change in FEV_1_ was 13 and 25 ml/yr over the first two years of treatment and fell by a mean of -28 to -65 ml/yr for years three to eight. DLco fell consistently for all eight years examined **(Figure 1B, C)**.

### Quantitative Proteomics of Human LAM Lung Tissue

We hypothesised that reduced responsiveness to rapamycin of patients with later stage disease may be consequence of progressive changes in the LAM cell microenvironment, providing rapamycin-insensitive pro-proliferative and pro-survival signals. To identify these signals, we investigated global proteome changes in explanted lung tissue from LAM patients undergoing lung transplantation (**Supplementary Figure 1**) relative to healthy controls, using a quantitative shotgun proteomics approach.

Lung tissue lysates were digested with trypsin, and proteins isotopically labelled; protein from healthy lung tissue was tagged by dimethylation with light formaldehyde (CH_2_O, +28 Da) and LAM lung tissue proteins were labelled with heavy formaldehyde (^13^CD_2_O, +34 Da) (**Figure 2A**); Data were analysed using MaxQuant [22,23] with a 1% False Discovery Rate (FDR) correction. 67 proteins were more than 1.3 fold upregulated in LAM lung relative to healthy lung **(Table 1).** To visualise these data, we used STRING [24] (https://string-db.org/) to investigate protein-protein interactions. STRING generated three distinct clusters of interacting proteins; using the inbuilt enrichment analysis these were identified as actin cytoskeleton, extracellular matrix, including type VI collagen, and glycolysis (**Figure 2B**). We also used Enrichr [17–19] to query Gene Ontology (GO) libraries producing similar results (**Figure 2C, Supplementary Table 1**). The enrichment of these proteins is consistent with accumulation of myofibroblasts in late stage LAM lung, as we have previously reported [5].

**Figure 2:**
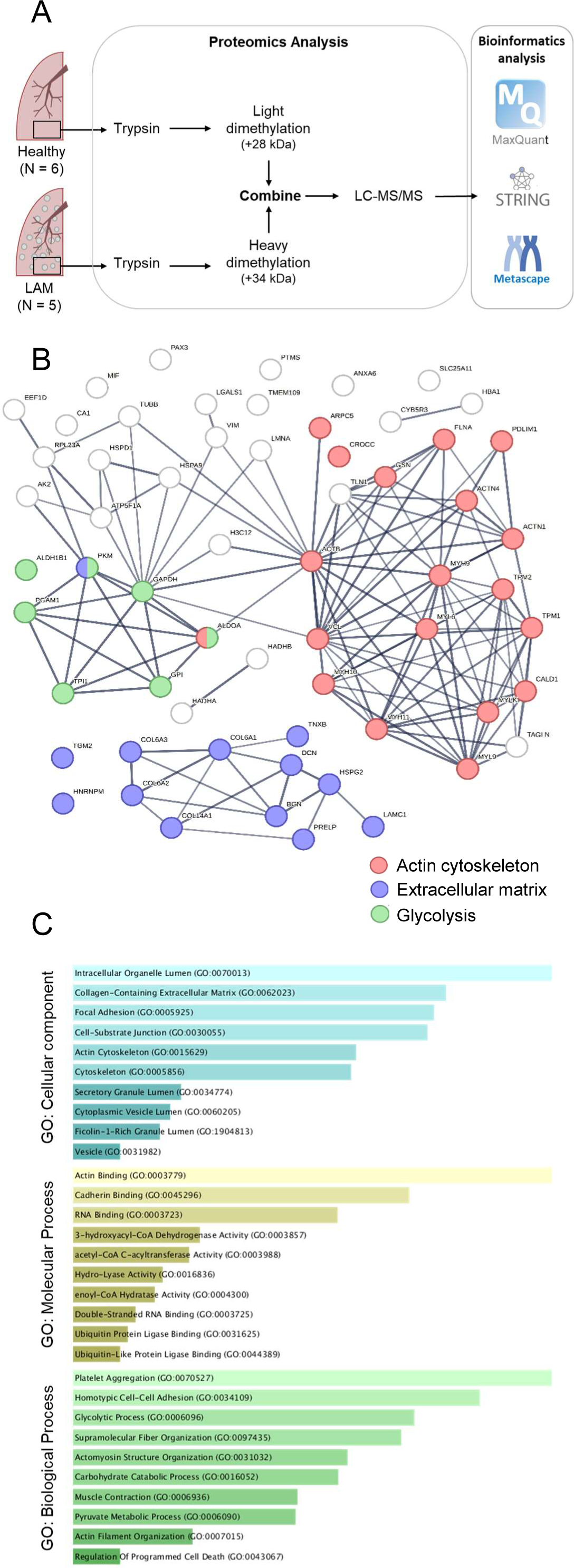
Quantitative proteomics of LAM lung tissue. (**A**) Outline of proteomics workflow and analysis. 136 proteins were identified as downregulated in LAM lung relative to healthy lung tissue, 113 as upregulated. A list of detected proteins is provided **Table 1** (**B**). Protein-protein interactions were mapped in STRING v12 with a list of 67 proteins which were upregulated >1.3 fold, with a confidence level of 0.7 (high) and 1% FDR. The resulting network had significantly more interactions than expected, with a Protein Protein Interaction (PPI) enrichment p-value < 1.0e^-16^. STRING pathway enrichment analyses identified the three major PPI clusters: actin cytoskeleton (red, Gene Ontology (GO) Cellular Compartment, p (FDR) = 1.89e^-13^), collagen containing extracellular matrix including Collagen VI (asterisks), (blue, p (FDR) = 1.88e^-08^) and glycolytic process (green, Gene Ontology Biological Process, p (FDR) = 2.13e^-05^). (**C**) Bar graph of enriched terms in GO libraries across input protein list, coloured by p values, for enrichments with adjusted p values <0.05. Refer to **Supplementary Table 1** for values.

**Table 1:**
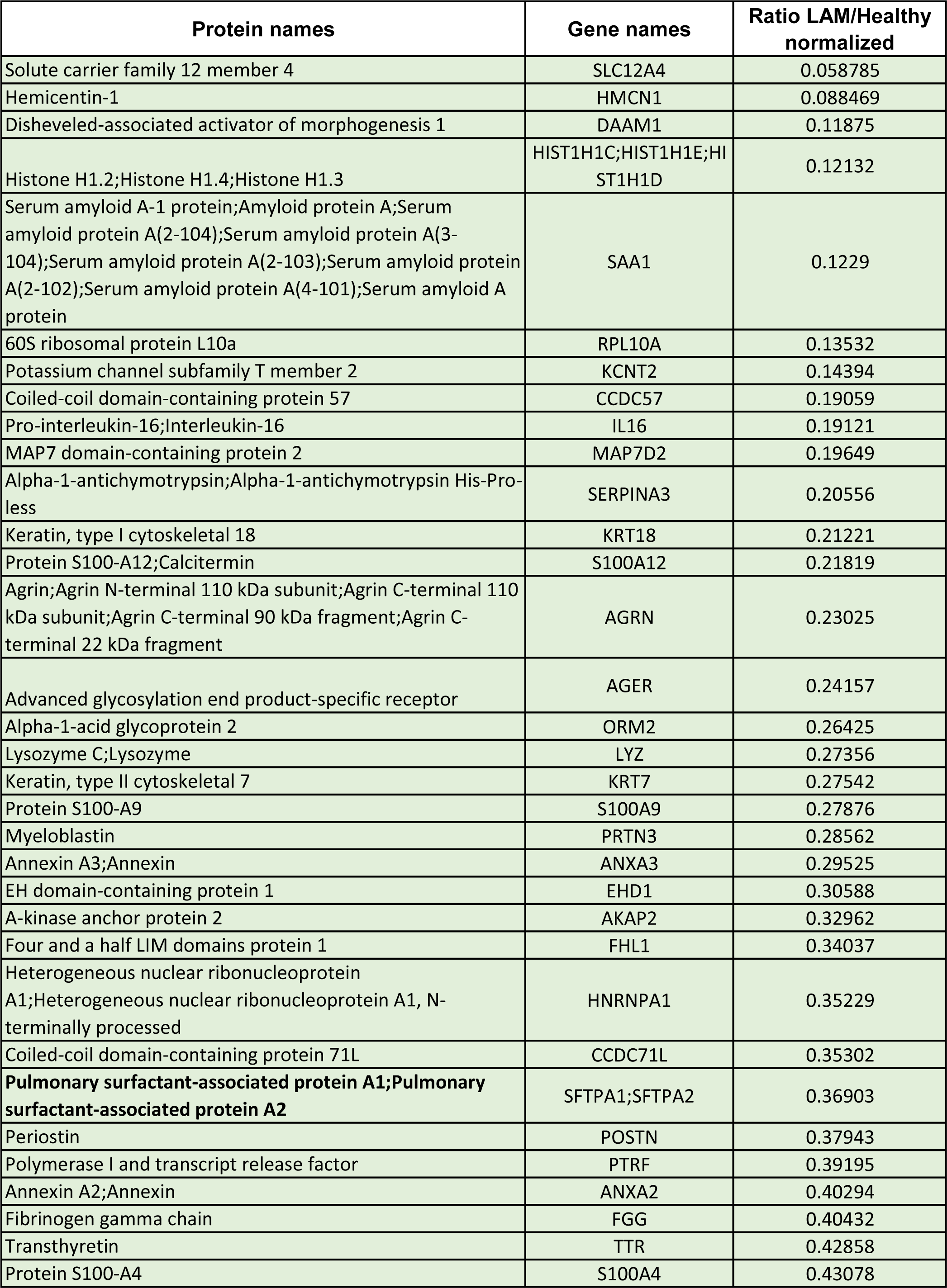
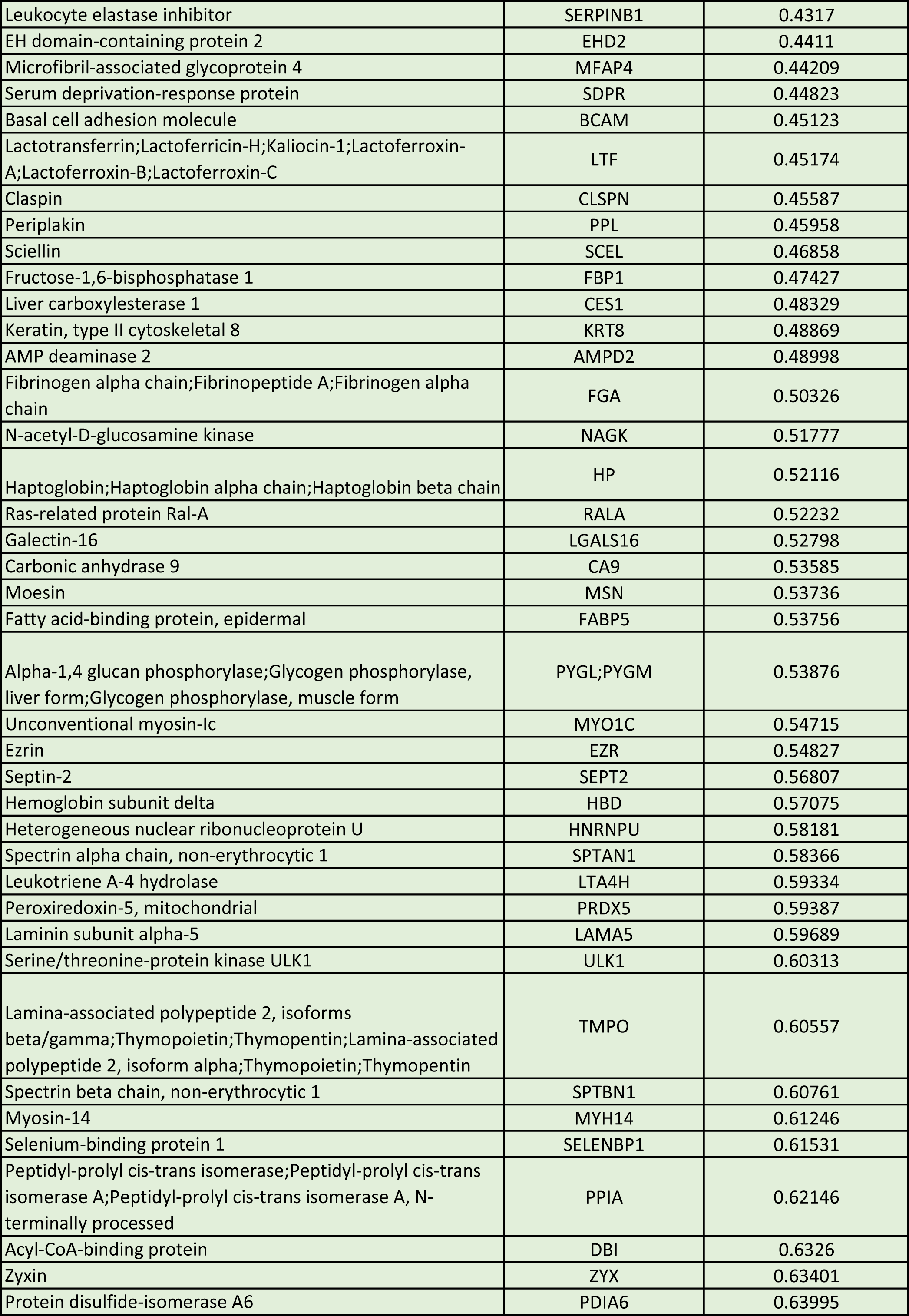
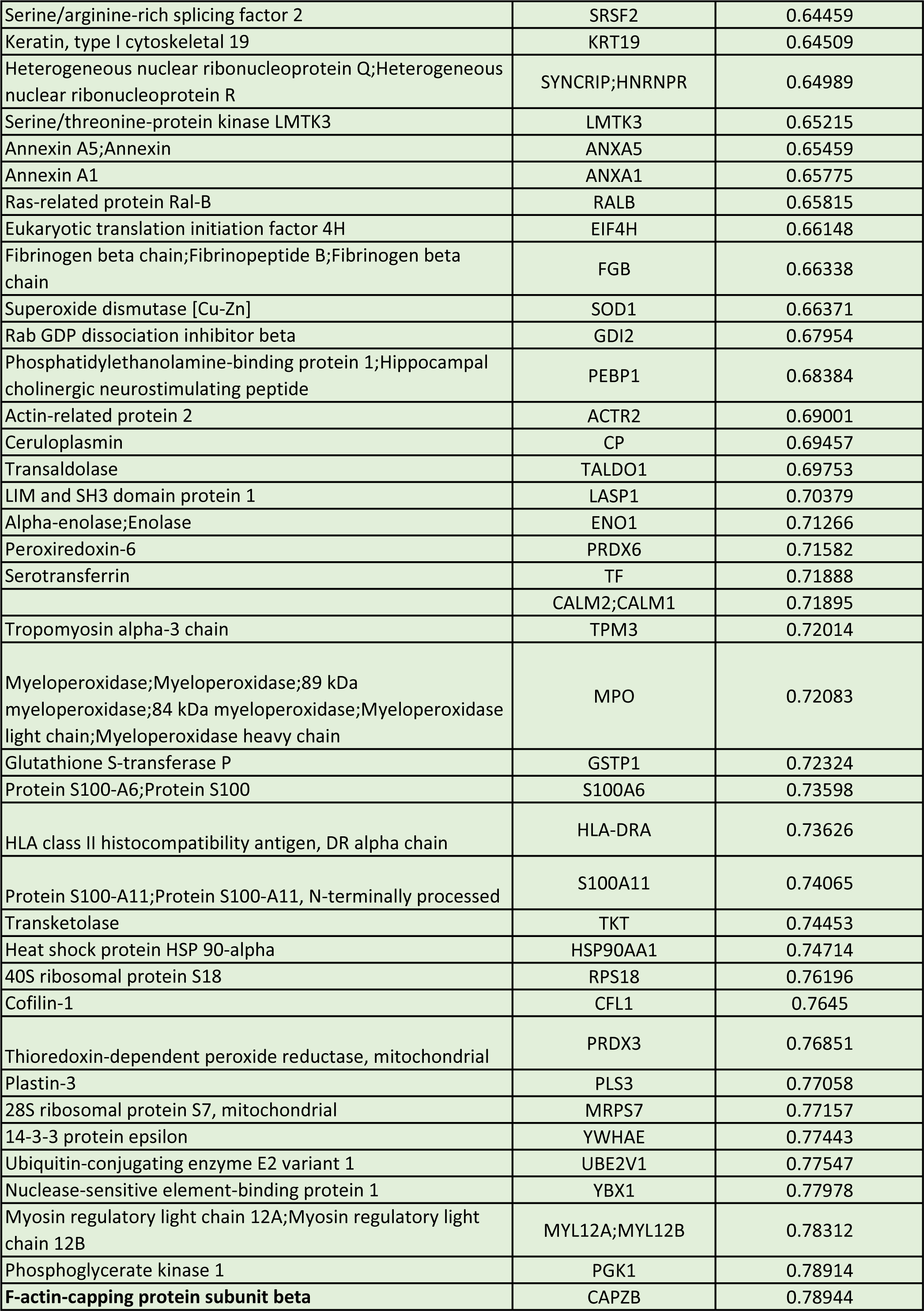
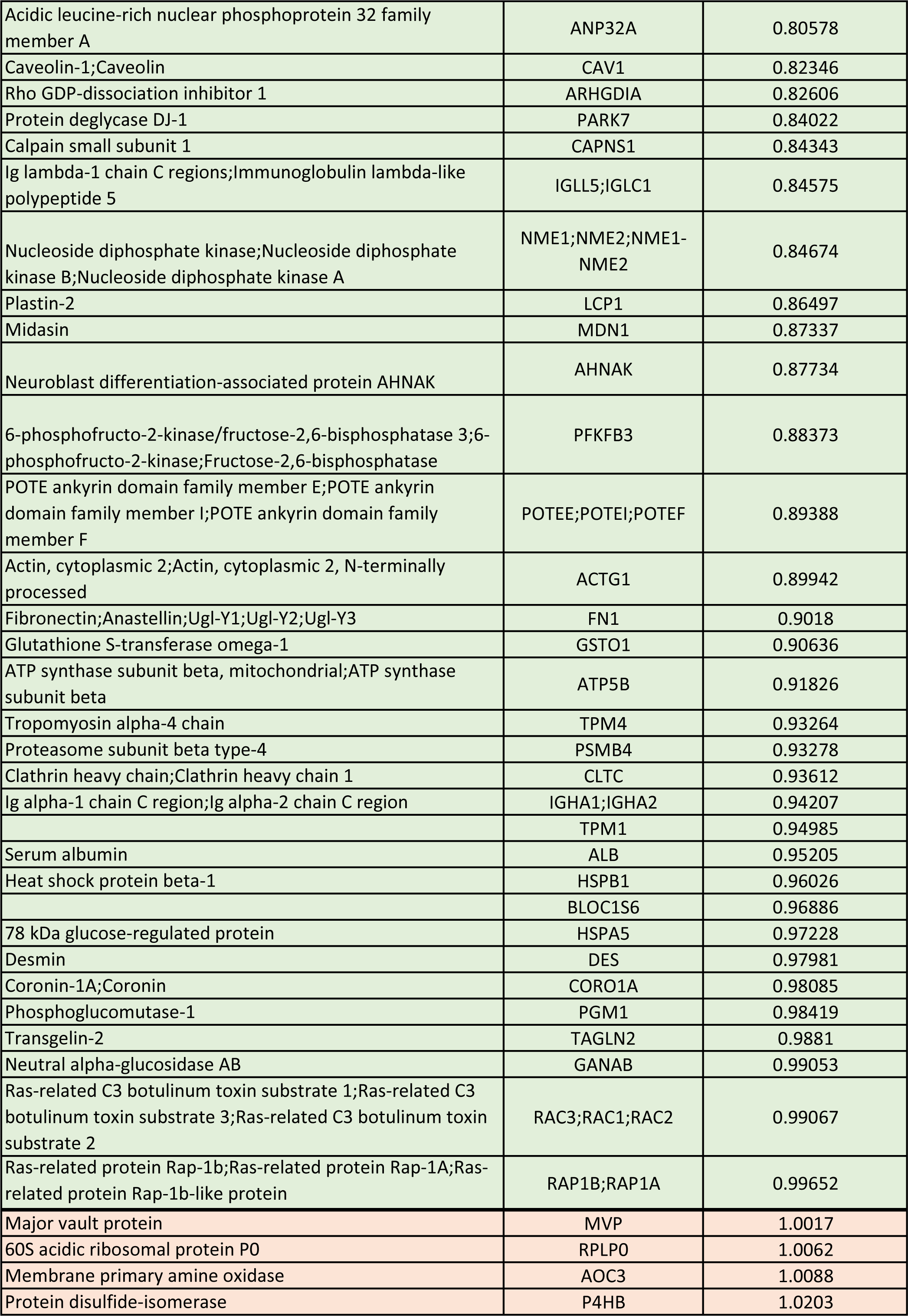
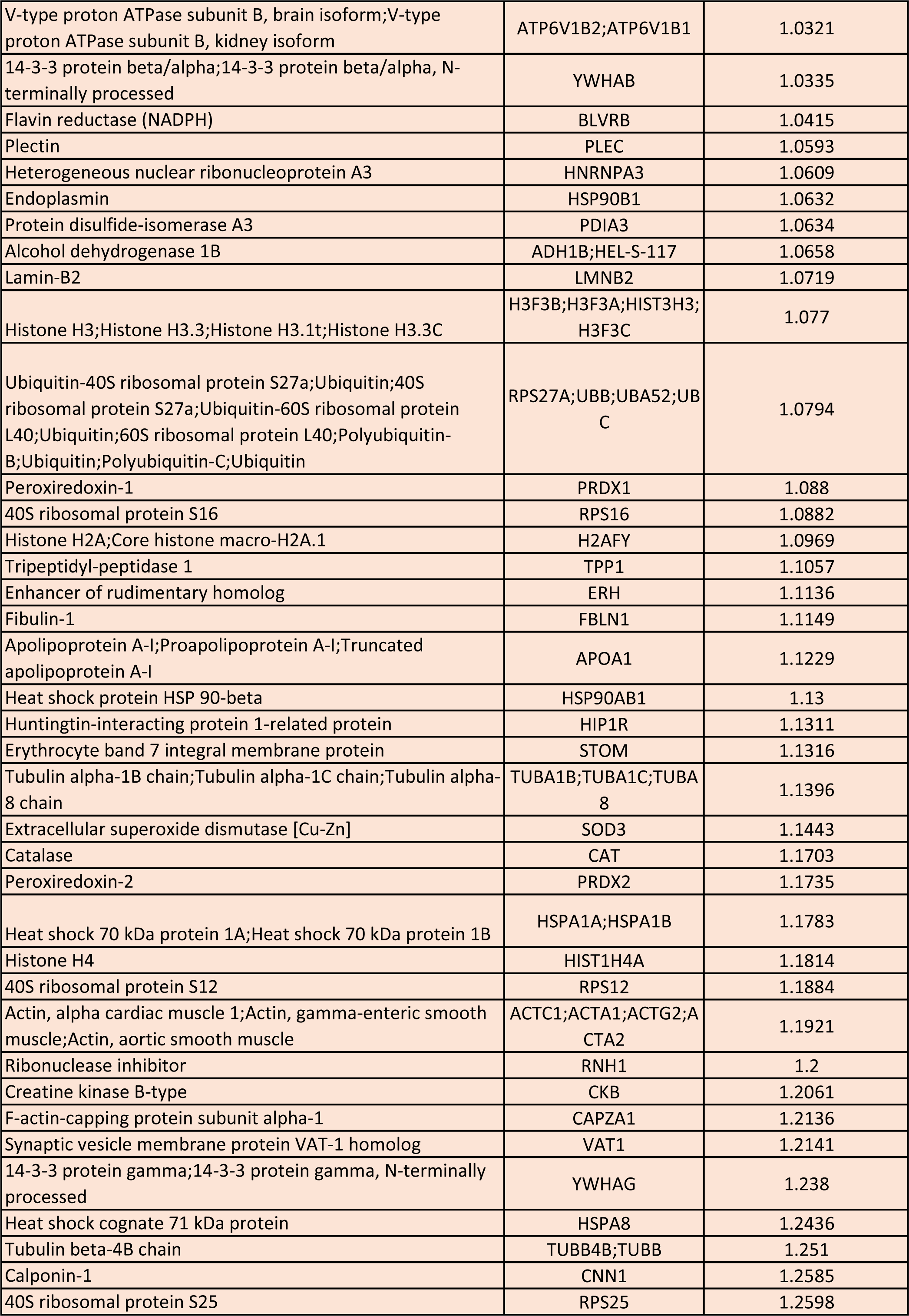
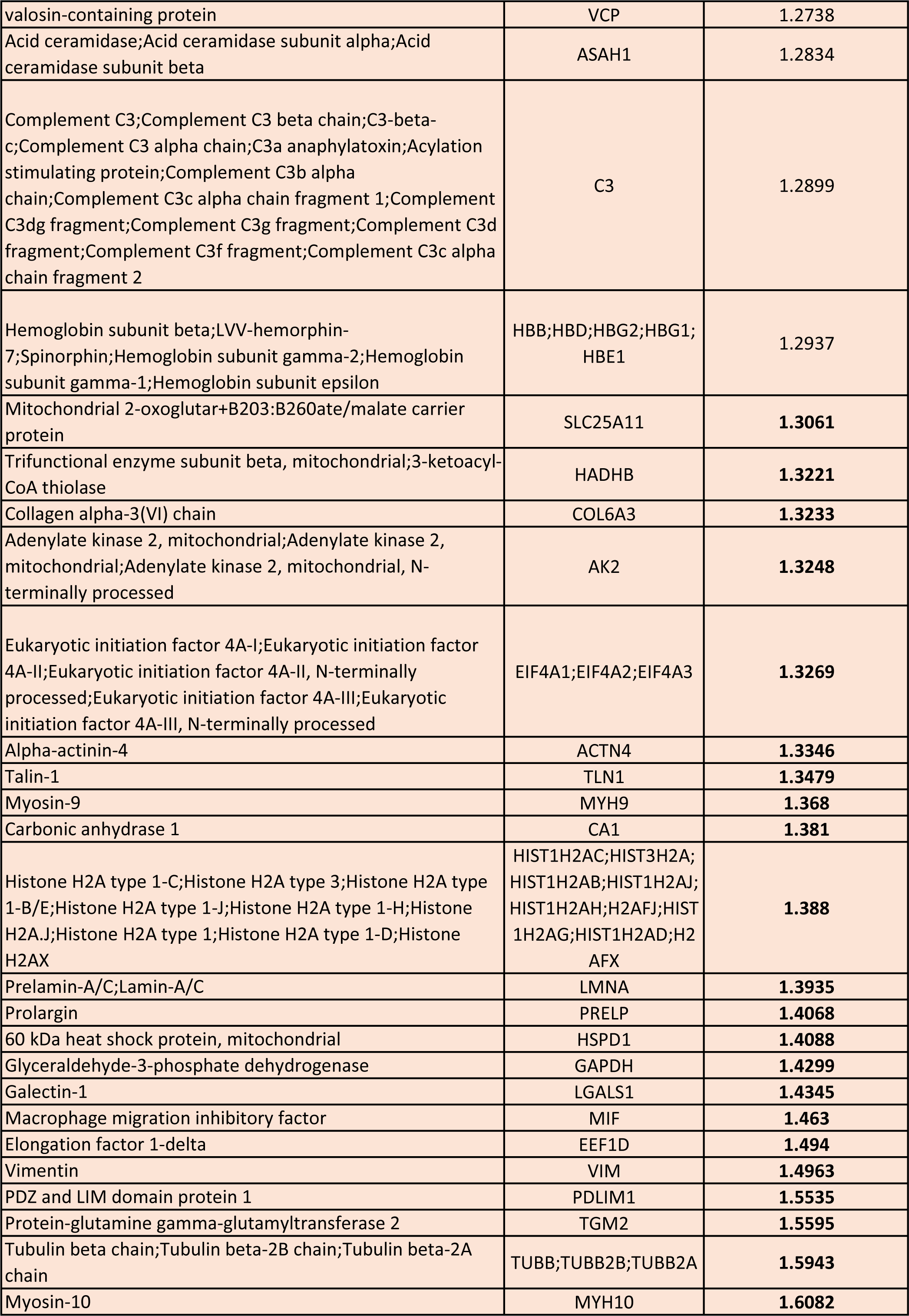
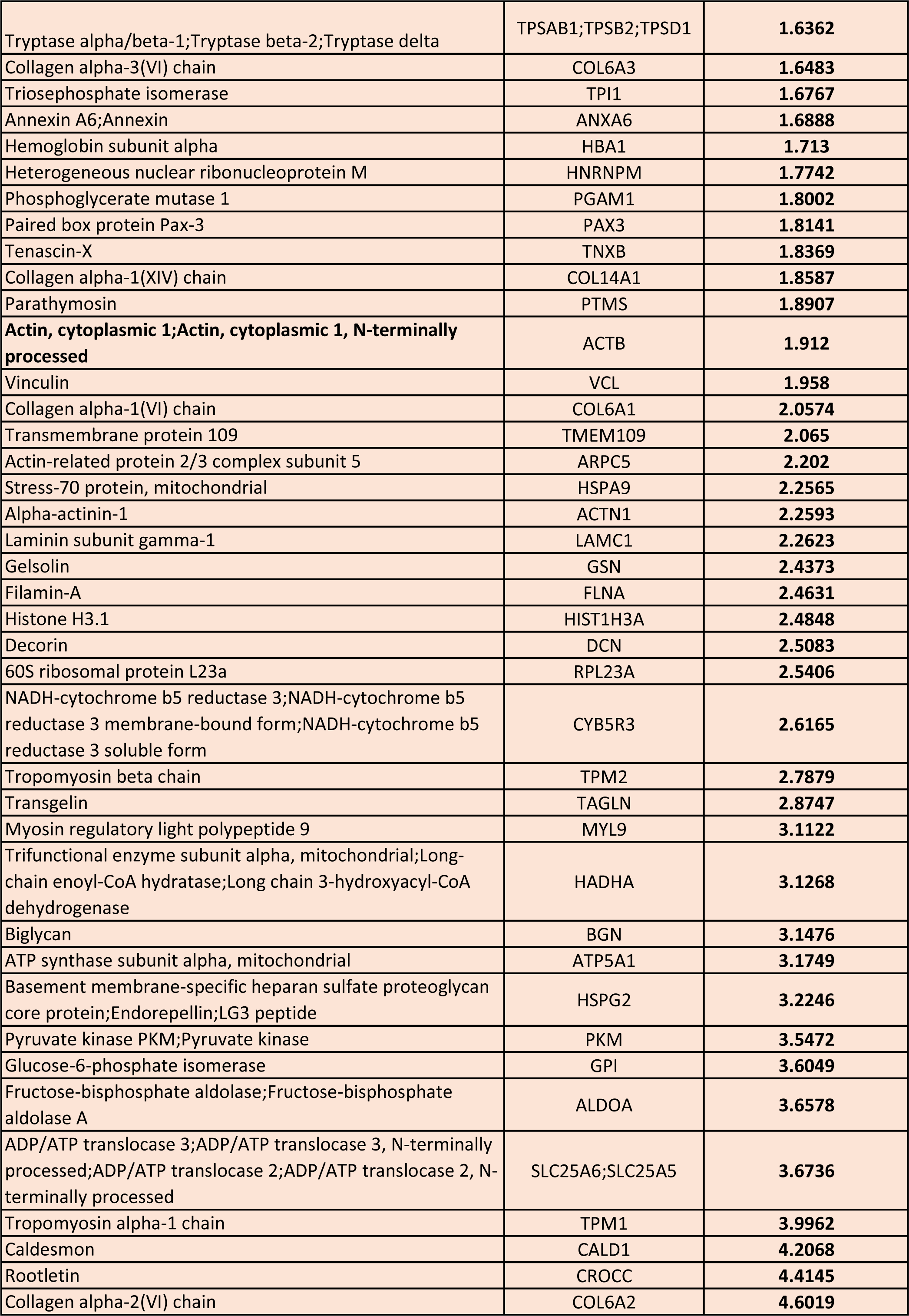
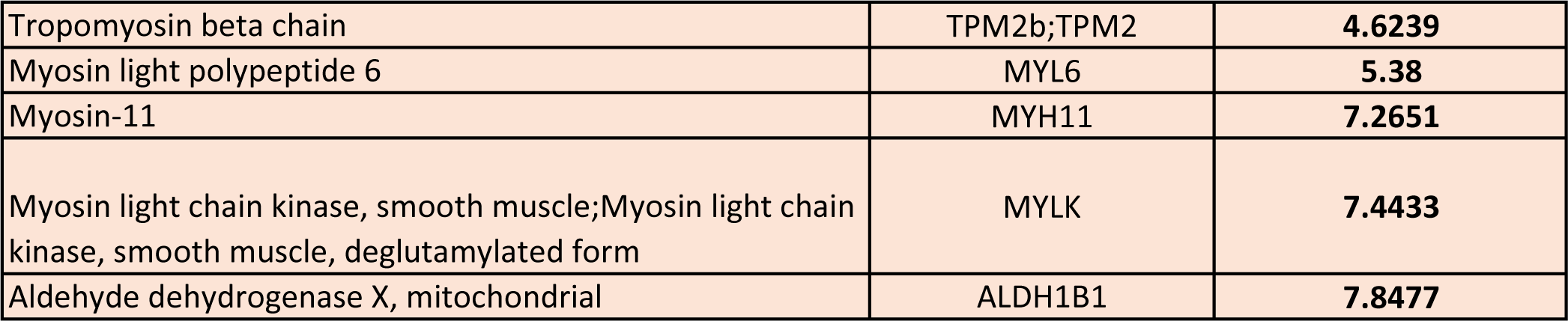
Proteins elevated (red) and decreased (green) in LAM patient lung tissue relative to tissue from healthy controls.

### Collagen expression in LAM patient samples

In order to determine whether ECM deposition was evident in LAM lung nodules we used picrosirius red (PSR) staining to visualise fibrillar collagens as a representative marker of ECM. Paraffin sections of lung tissue from 24 patients with linked clinical information and from three explanted lungs from patients with LAM were stained with PSR and imaged **(Figure 3A)**. Collagen staining was evident in nodules by this method; to quantify the degree of collagen deposition, the area of PSR staining relative to the area of a nodule was calculated for multiple nodules from each patient. We investigated the relationship between degree of collagen deposition and the lung function parameters % predicted FEV_1_ (ppFEV_1_) and DL_co_, (ppDLco) and disease duration, but found no significant correlation. **(Figure 3B)**.

**Figure 3:**
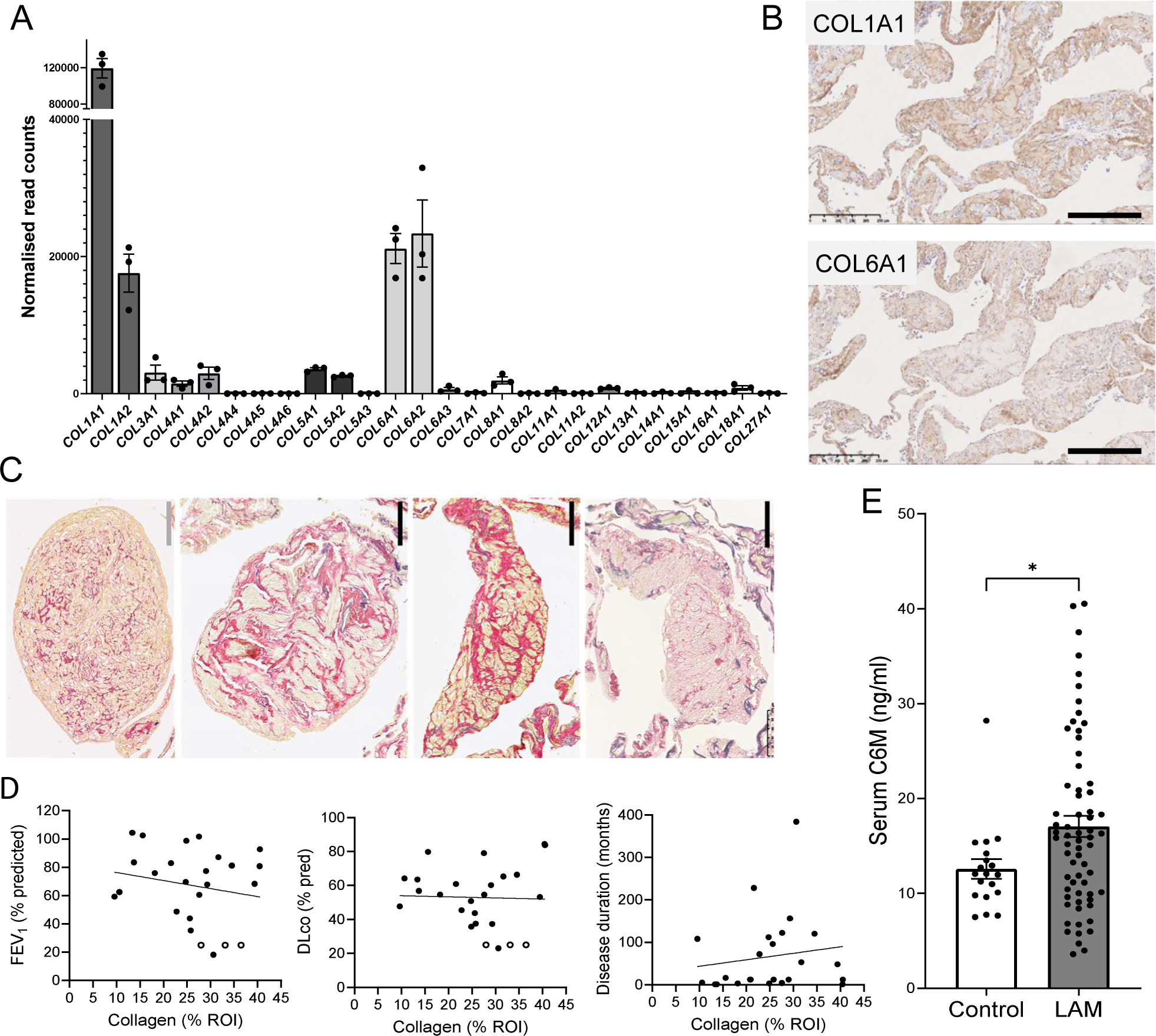
Collagen expression in LAM. (**A**) Expression of collagen genes by primary LAM Associated Fibroblasts in tissue culture measured by RNA sequencing reveals *COL1A1*, *COL1A2*, *COL6A1* and *COL6A2* as the most highly expressed transcripts. (**B**) Both Collagen alpha-1(I) chain (COL1A1) and Collagen alpha-1(VI) chain (COL6A1) can be detected in areas of the lung affected by LAM by immunohistochemistry (scale bar = 250 µm). (**C**) Picrosirius red staining (PSR) reveals varying degrees of fibrillar collagen deposition with LAM lesions (scale bars: grey = 50 µm, black = 100 µm). (**D**) Quantification of collagen deposition in lesions (area of PSR staining/area of lesion) No significant correlation was detected between collagen deposition and ppFEV_1_ (R^2^ = 0.037, p = 0.356), ppDL_CO_ (R^2^ = 0.001, p = 0.88) or disease duration (R^2^ = 0.021, p = 0.49). Open circles represent imputed data for explanted lung tissue from patients with end-stage disease. (**E**) Serum levels of an MMP-derived neoepitope of collagen alpha-1(VI), C6M, are significantly elevated in LAM patients relative to healthy controls (p = 0.0427, Mann-Whitney test).

The three main chains (A1, A2 and A3) of the mature type VI collagen molecule were noted in the ECM cluster of proteins upregulated in LAM lung tissue shotgun proteome relative to healthy lung **(Figure 2B)**. In LAM patient serum an MMP-derived neoepitope of Collagen alpha-1(VI) chain, C6M, was significantly elevated (p = 0.0427) relative to healthy control serum (mean of 66 LAM patients = 17.06, 95% CI = 14.82-19.28, mean of 19 healthy controls = 12.57, 95% CI = 10.37-14.77) (**Figure 3C**). By RNA sequencing, expression of *COL6A1*, *COL6A2* and *COL6A3* was found in cultured primary LAFs, although *COL1A1* was the most abundantly transcribed collagen gene (**Figure 3D**). Both Collagen alpha-1(VI) chain and Collagen alpha-1(I) chain were detected in LAM lesions by immunohistochemistry (**Figure 3E**)

### Fibroblast-deposited ECM and cell proliferation

The supportive role of ECM in promoting growth of cells in culture is well-documented [25] and we hypothesised that contact with fibroblast-deposited ECM could stimulate LAM cell proliferation *in vivo*. To test this *in vitro* we prepared cell culture plates coated with LAF ECM by decellularisation of confluent LAF culture plates from three LAF donors, and grew LAM patient-derived 621-101 cells, a model cell line for LAM [20], on this substrate and on standard tissue culture plastic for five days, in culture medium containing low (1%) serum. 621-101 cells grown on LAF ECM were significantly more proliferative then those grown on plastic when assayed by cell counts (p = 0.0018) **(Figure 4A, B).** We also used BrdU incorporation, resazurin reduction and MTT reduction to investigate the effect of LAF ECM on proliferation over 72 hours. These approaches produced comparable results, showing a significant increase in proliferation and/or viability of cells grown on LAF ECM relative to plastic (**Figure 4C**). To determine whether the observed increase in proliferation of 621-101 cells grown on ECM was specific to ECM deposited by LAM-derived fibroblasts, we also generated culture plates coated with ECM deposited by fibroblasts isolated from normal lung tissue (Normal Human Lung Fibroblasts, or NHLFs). NHLF ECM and LAF ECM were not significantly different in their ability to support increased proliferation (not shown).

**Figure 4:**
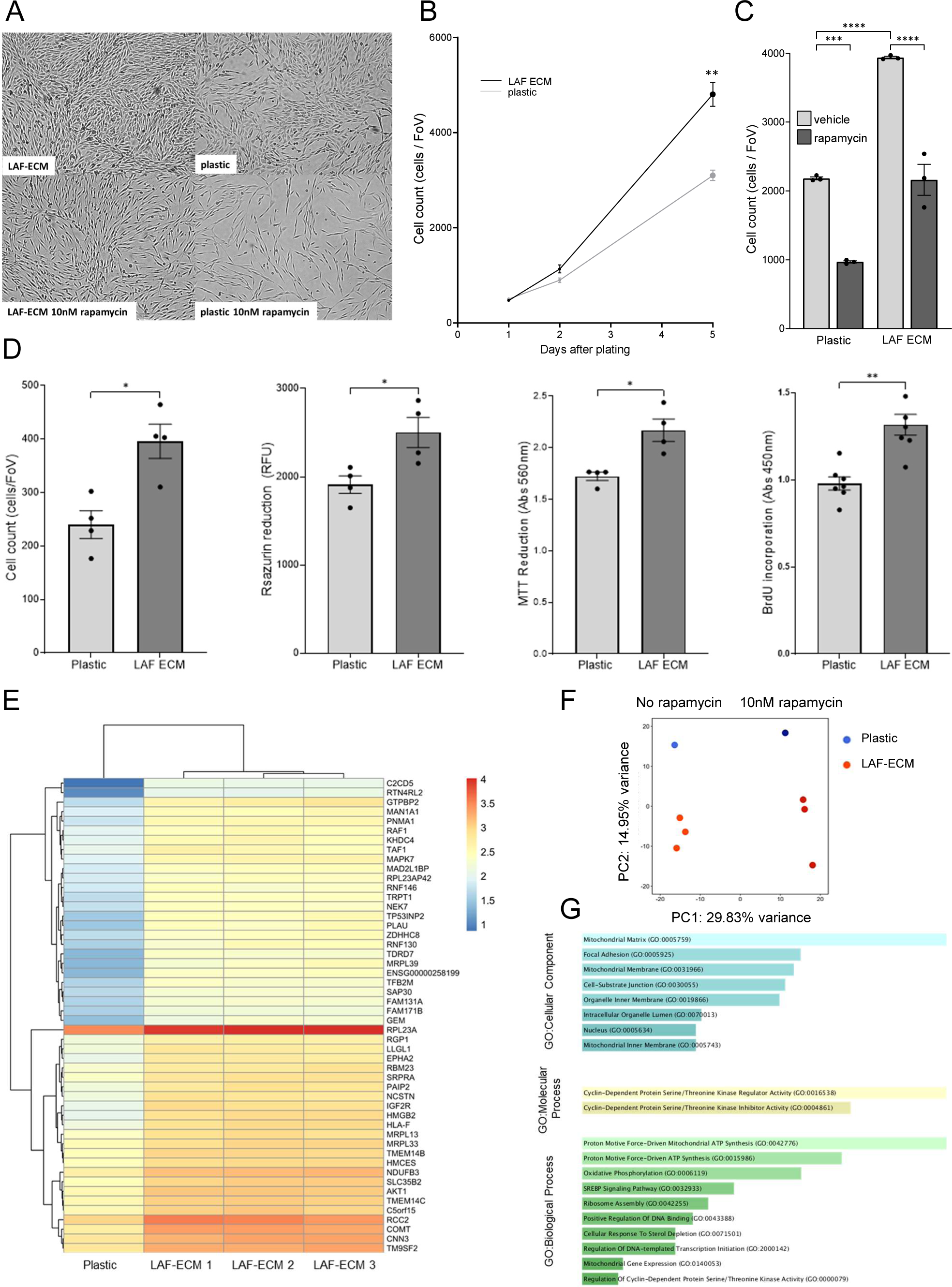
Effect of decellularised LAF ECM on proliferation of 621-101 cells. Patient-derived TSC2^-/-^ 621-101 cells, a model LAM cell line, proliferate more rapidly on plates coated with decellularised LAF-deposited extracellular matrix (LAF ECM) than on tissue culture treated plastic. (**A**) Brightfield image (10X), (**B**) cell counts of cells grown for five days in tissue culture medium containing 1% FCS on plastic or LAF ECM. This growth advantage is maintained in the presence of 10 nM rapamycin (**C**) and is consistent when measured by a range of proliferation/viability assays: cell counts, ‘viability’ (resazurin and MTT reduction) and cell cycle progression. * p<0.05, **p<0.01, ***p<0.001, ****p<0.0001 by ANOVA and t-test. (**D**). RNA sequencing was performed on 621-101 cells grown for five days on plastic or LAF-ECM, in medium containing 1% FCS, in the presence and absence of 10 nM rapamycin (n=3 LAF matrix donors). Growth on LA -ECM elicited changes in gene expression in 621-101 cells. The heatmap (**E**) shows the top 50 differentially expressed genes obtained by sorting the absolute value of fold change. (**F**) Principal component analysis (PCA) of all samples. Principal component (PC) 1 divides the samples by rapamycin treatment, PC2 divides the samples by plastic/ECM growth substrate. (**G**) Pathway analysis of genes upregulated on LAF ECM with log2FC>0.5, padj<0.05 revealed enrichment of pathways involving mitochondrial activity, cyclin-dependent kinase activity, and cell-substrate interaction (see **Supplementary Table 5** for p values).

Rapamycin has a significant cytostatic effect on 621-101 cells [26]. To determine whether LAF ECM was able to maintain 621-101 cell proliferation in the presence of rapamycin we plated these cells on plastic or LAF ECM and grew them in the presence or absence of 10 nM rapamycin for five days. Although rapamycin significantly decreased proliferation on both substrates (p = 0.0003 for cells on plastic, p<0.0001 for cells on LAF ECM), there were significantly more cells after five days on LAF ECM relative to plastic (p = 0.0003) (**Figure 4D**)

### Transcriptional response of cells to LAF ECM

To identify gene expression changes in 621-101 cells grown on LAF ECM relative to standard tissue culture plastic, and investigate the effect of rapamycin on transcription in cells grown on these substrates, RNA sequencing was performed on cells grown on plastic (control) or LAF ECM derived from three LAF donors, for five days, in the presence and absence of 10 nM rapamycin (**Figure 4E, F; Supplementary Tables 2, 3, 4)**.

We identified 402 genes with upregulated expression in cells grown on LAF ECM and 53 genes with downregulated expression, using log2 fold change (log2FC)>0.5 or <-0.5 and adjusted p value (p_adj_)<0.1. (**Table 2**). To identify enriched pathways amongst upregulated genes we used Enrichr [17–19] to interrogate GO libraries, which revealed significant upregulation of pathways driving metabolism, transcription and cell cycle control, consistent with increased cell proliferation (**Figure 4G, Supplementary Table 5**).

**Table 2:**
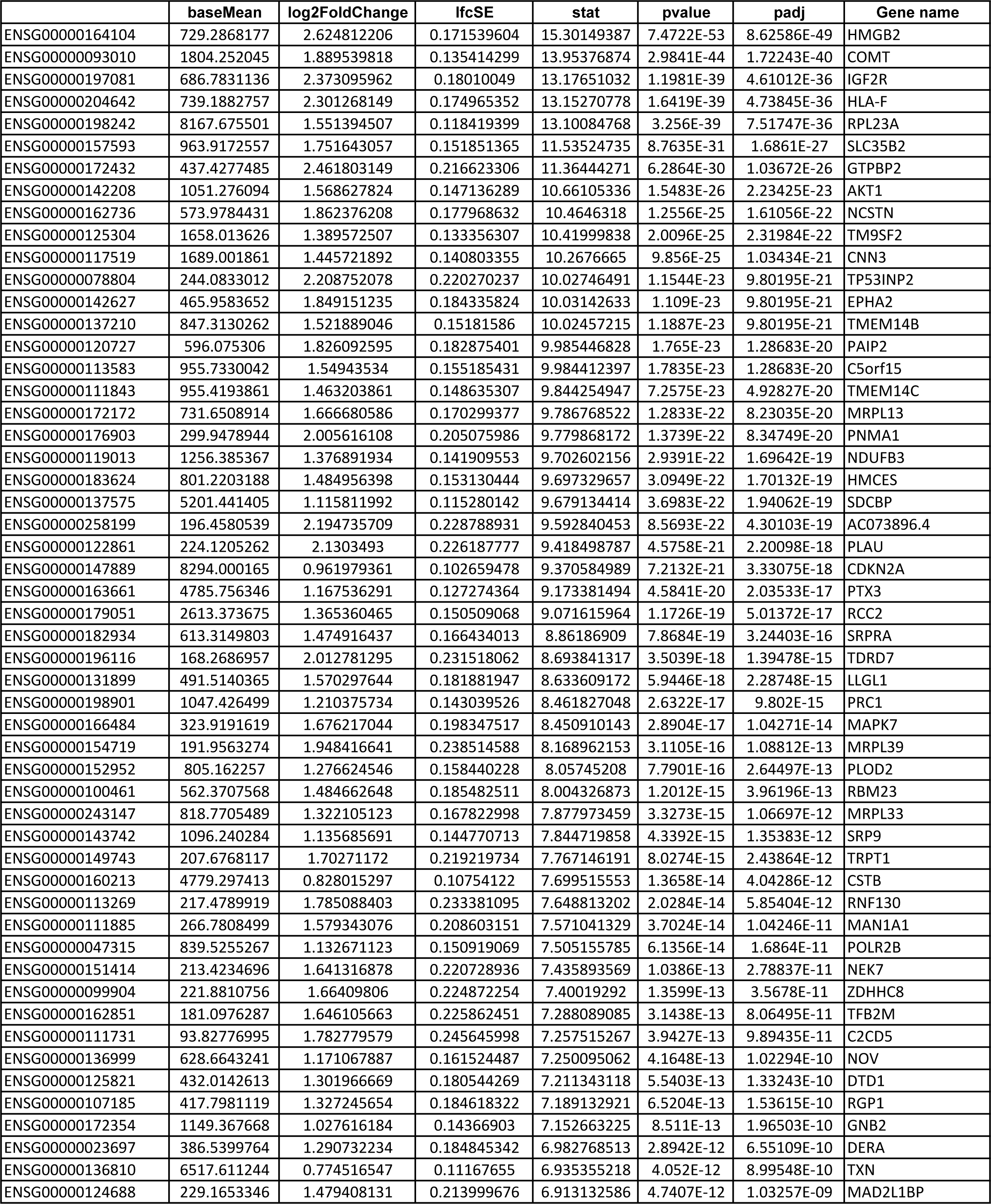
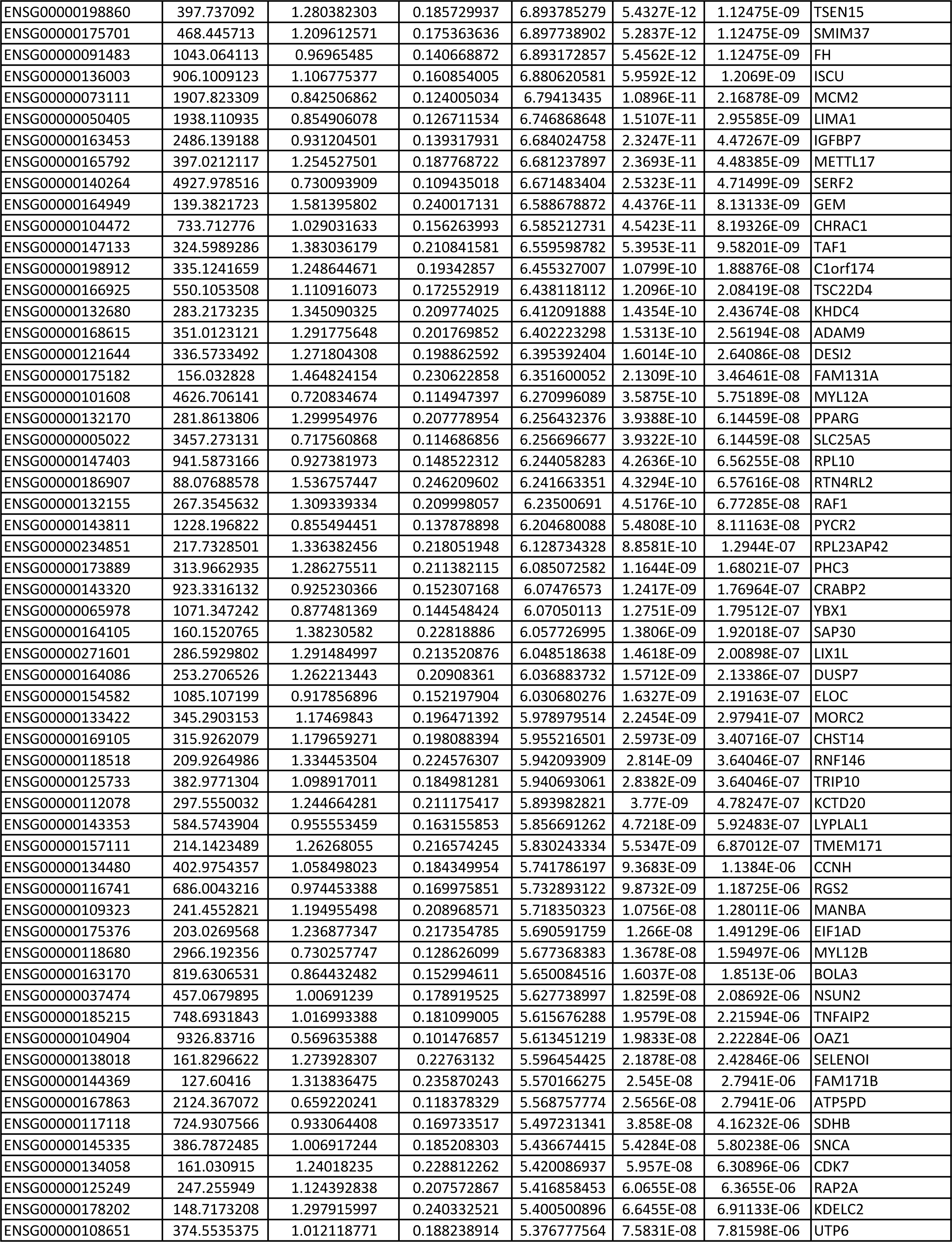
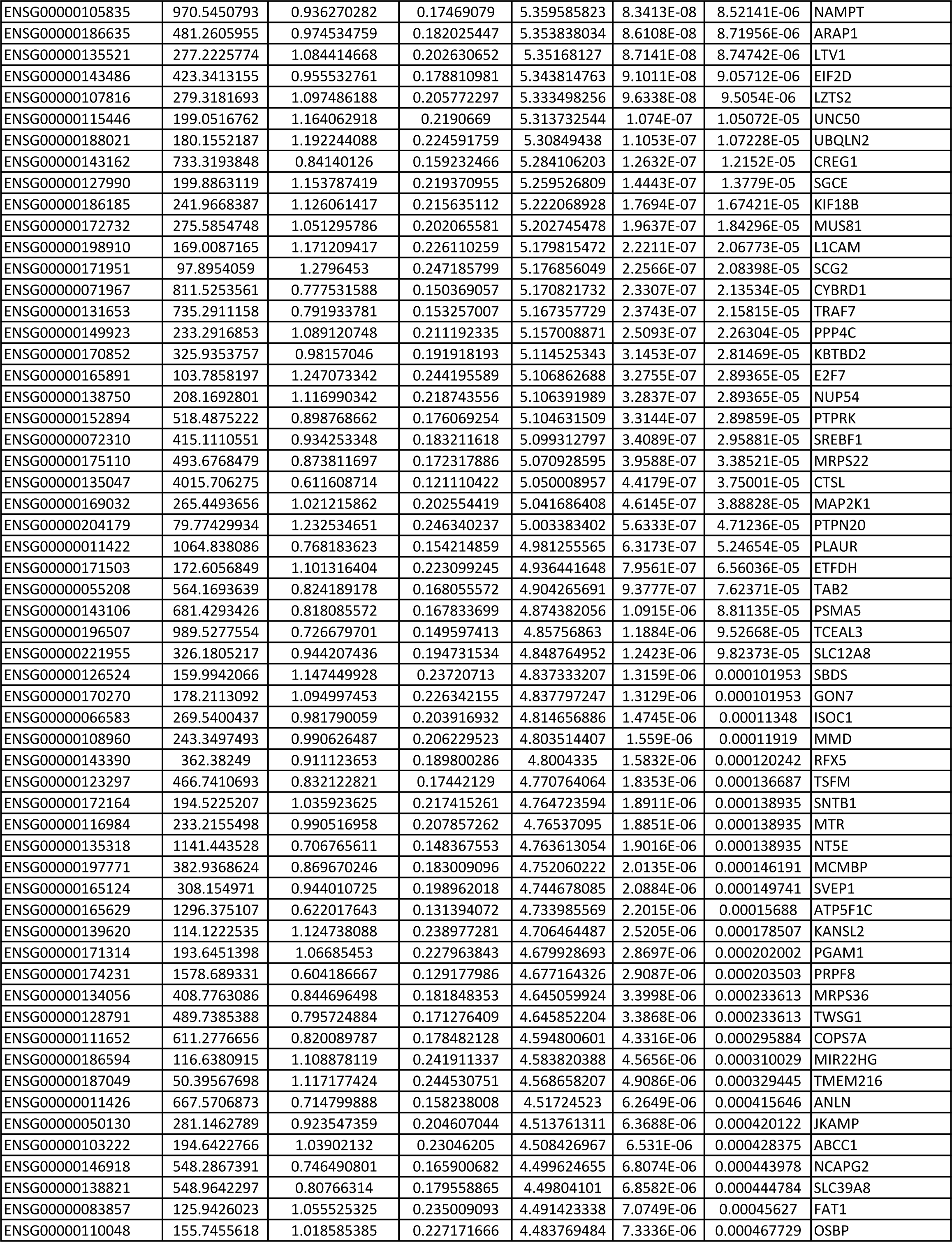
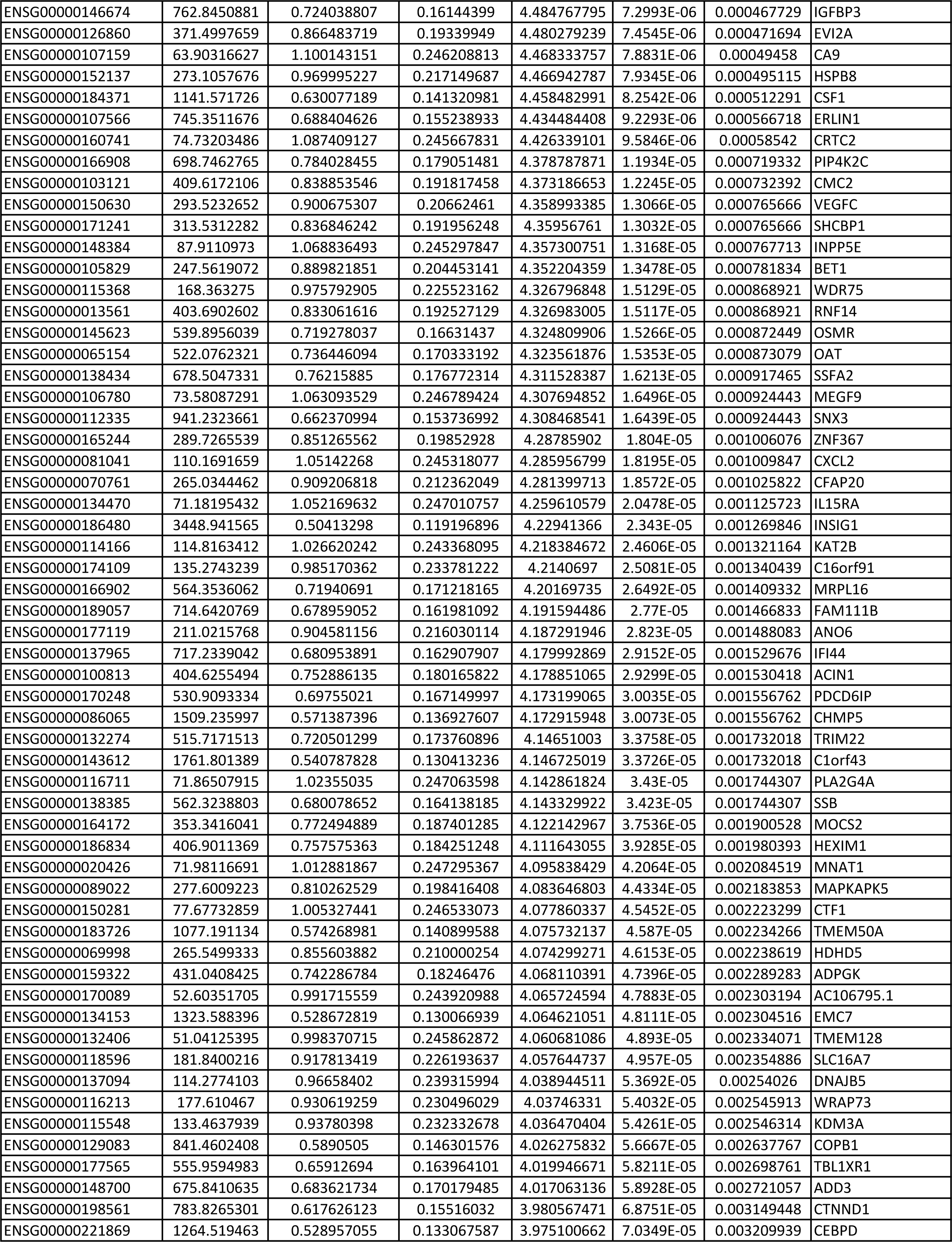
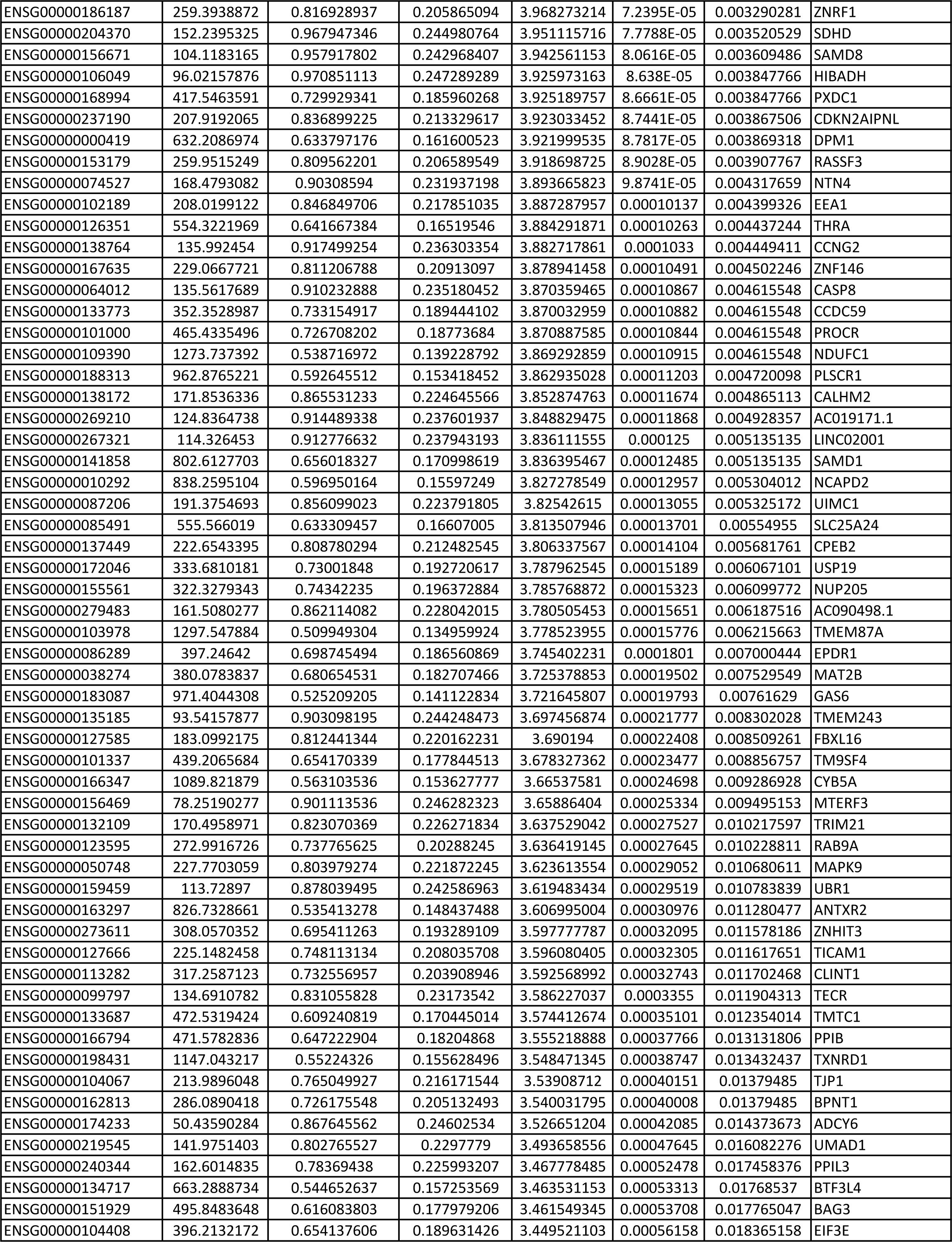
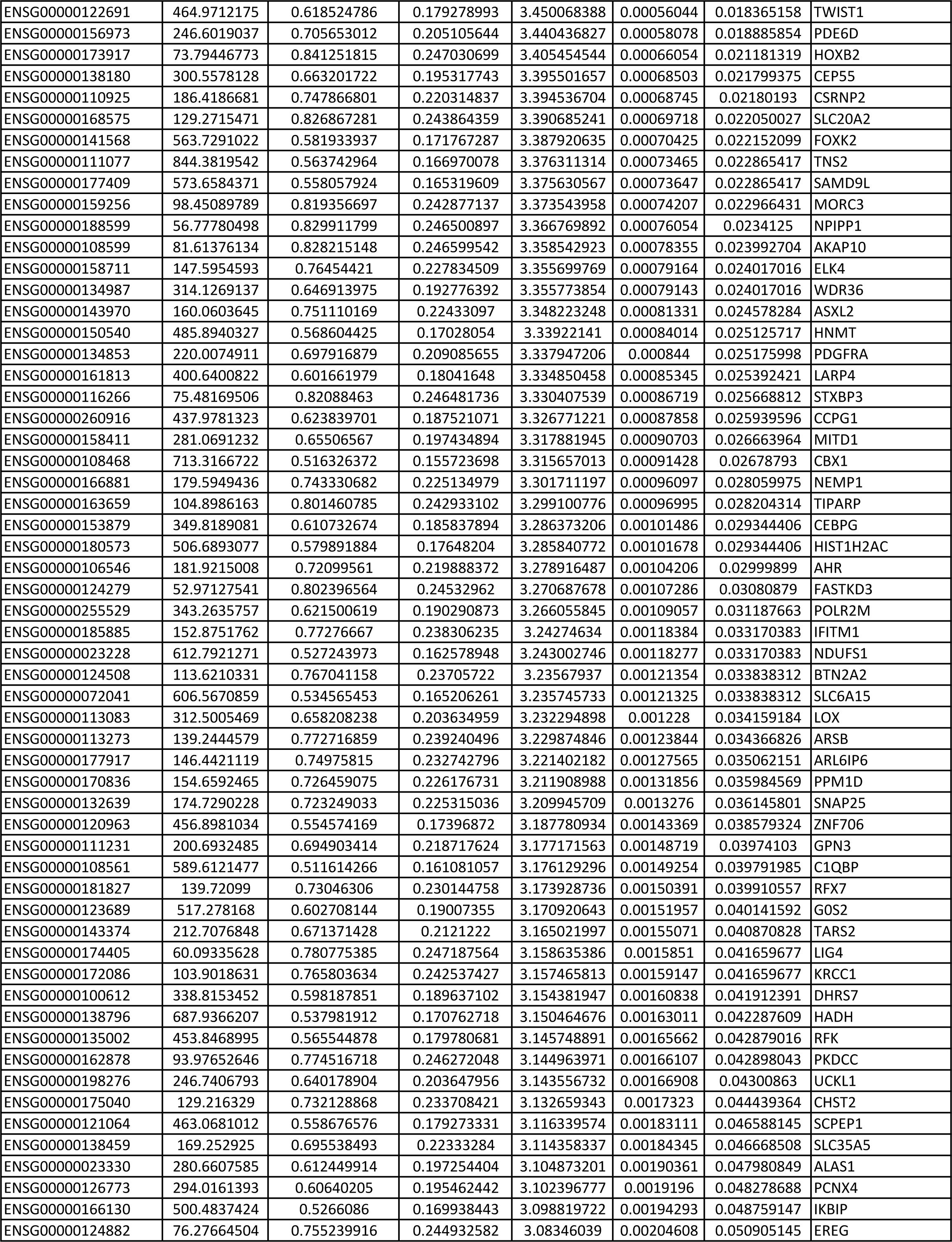
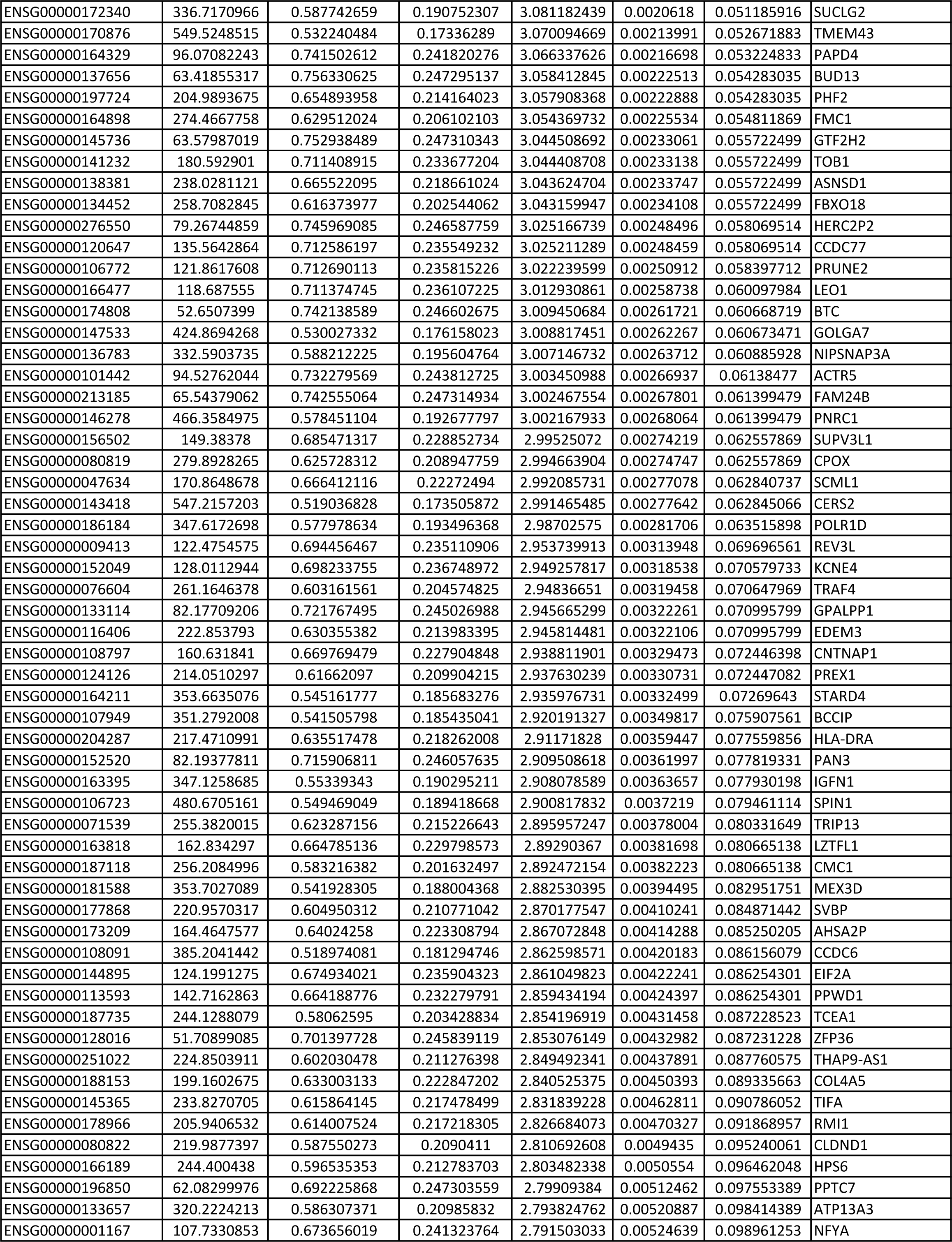
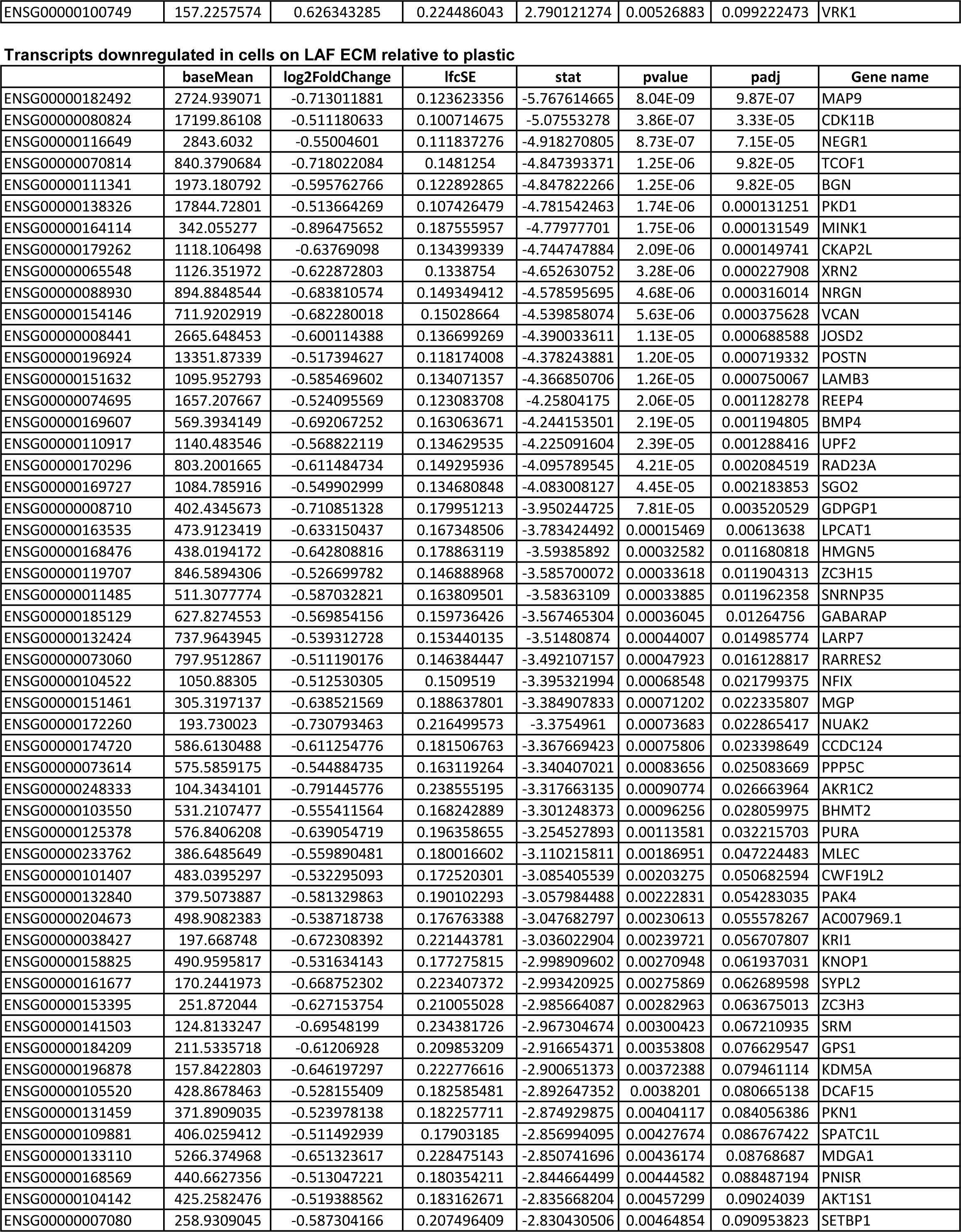
Transcripts upregulated and downregulated ((log2FC)>0.5 or <-0.5 and adjusted p value (padj)<0.1) in 621-101 cells grown on LAF-deposited ECM relative to cells grown on plastic. For full data see **Supplementary Tables**.

We interrogated these data to identify genes encoding druggable components of pro-proliferative pathways. We also determined whether the expression of these genes was sensitive to inhibition by rapamycin, since we were seeking pathways that could drive proliferation in the presence of rapamycin. Genes identified which fulfilled these criteria included: *Cyclin-Dependent Kinase 7* (*CDK7*), a member of the both the Cyclin Activating Kinase (CAK) complex and the core transcription factor TFIIH, *Growth Arrest Specific 6* (*GAS6*), and both *Plasminogen Activator, Urokinase* (*PLAU*), encoding the serine protease urokinase, and its receptor *Urokinase-type Plasminogen Activator Receptor* (*PLAUR,* also known as uPAR) (**Figure 5A, 5B, 5C**). The expression of the additional two components of the CAK complex, *MNAT1* (MAT1) and *CCNH* (Cyclin H) were also seen to be significantly upregulated in cells grown on LAF ECM. GAS6 is a ligand for members of the TAM (Tyro3, AXL, MerTK) family of receptor tyrosine kinases; of these receptors, *AXL* is the most highly expressed in 621-101 cells (**Supplementary Table 2**), and although its expression is not elevated in cells grown on LAF ECM relative to plastic, it is upregulated when cells are treated with rapamycin (log2FC = 0.916, p_adj_ = 4.05E^-25^, **Supplementary Table 3**) and so potentially contributes to rapamycin-insensitive cell proliferation. We used immunohistochemistry to verify the expression of the cognate proteins of these genes in LAM lung tissue; all were seen to be present in LAM lesions, which were identified by reactivity to the anti-melanoma antibody PNL2 [27] **(Figure 5C**).

**Figure 5:**
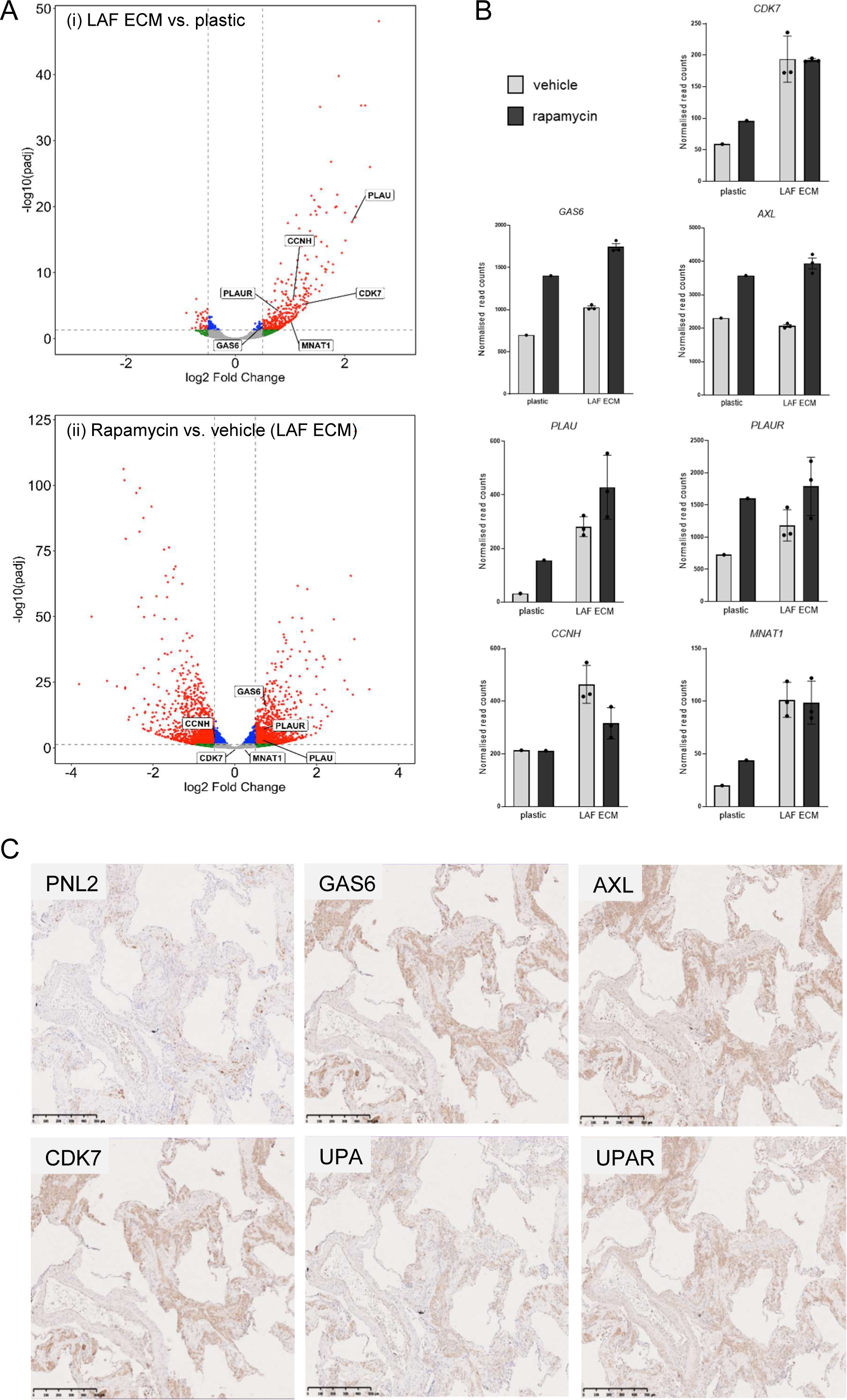
Identification of potential mediators of LAF-ECM driven cell proliferation. (**A**) Volcano plot of differentially expressed genes identified between 621-101 cells grown on tissue culture treated plastic and LAF ECM (i), and between cells grown on LAF ECM in the presence and absence of 10 nM rapamycin (ii). Genes selected for further investigation (*CDK7*, *PLAU*, *PLAUR* and *GAS6*) were found to be upregulated in cells grown on LAF ECM relative to plastic, with log2FC>0.5, padj<0.05, and unaffected or upregulated by rapamycin treatment. *MNAT1* and *CCNH* encode the additional components of the CAK complex, MAT1 and Cyclin H, while AXL is the likely receptor for GAS6. (**B**) Graphical representation of normalised read counts for genes of interest (refer to **Supplementary Table 2B**). *AXL* expression is not elevated in cells grown on LAF ECM, but is elevated by rapamycin treatment. (**C**) Protein expression was demonstrated by immunohistochemistry, LAM cells identified by reactivity to the anti-melanoma antibody PNL2. Scale bar = 500 µm, representative of six LAM tissue donors.

### Effect of pathway inhibitors on cell proliferation

To investigate whether CDK7, PLAUR/PLAU and GAS6/AXL pathways contribute to the ability of ECM to stimulate LAM cell proliferation we used inhibitors of these pathways to determine whether blocking their activity could attenuate ECM-driven proliferation. To target CDK7 we used the ATP competitive inhibitor Samuraciclib/CT7001 [28], for GAS6/AXL we used the multikinase inhibitor Dubermatinib/TP-0903 [29] and the TAM kinase specific inhibitor LDC1267 [30], and for PLAUR/PLAU we used IPR-803, a small molecule inhibitor of the PLAU/PLAUR interaction [31].

621-101 cells grown on plastic or LAF ECM were treated with these inhibitors and growth assayed by resazurin reduction. Samuraciclib, Dubermatinib and IPR-803 significantly decreased proliferation of 621-101 cells grown on both plastic and LAF ECM (**Figure 6A**). In combination with rapamycin, these drugs were able to reduce proliferation to a greater extent than 10 nM rapamycin alone (**Figure 6B**). LDC1267 showed no anti-proliferative effect (not shown).

**Figure 6:**
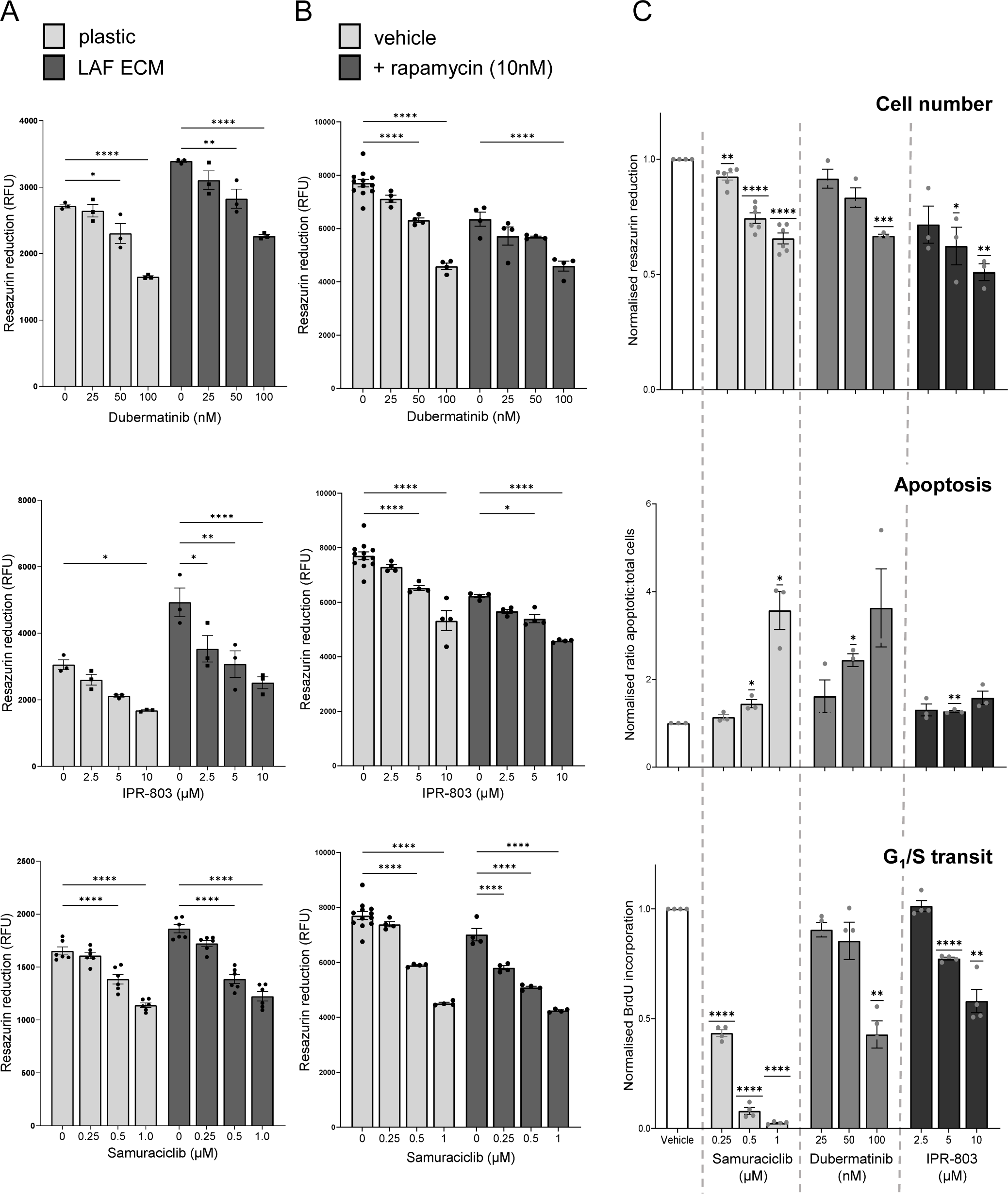
Effect of inhibitors of selected pathways on 621-101 proliferation. (**A**) Dubermatinib (GAS6/AXL), IPR-803 (PLAU/PLAUR) and Samuraciclib (CDK7) significantly inhibit proliferation of 621-101 cells grown on both plastic and LAF ECM substrates (statistical analysis by ANOVA, * p<0.05, **p<0.01, ***p<0.001, ****p<0.0001. (**B**) In the presence of 10 nM rapamycin, Dubermatinib, IPR-803 and Samuraciclib reduce viability of cells grown on LAF ECM significantly further than rapamycin alone. (**C**). Doses of these inhibitors which cause similar decreases in overall viability when cells are grown on LAF ECM, measured by resazurin reduction, achieve this by different mechanisms. Apoptosis was assayed by *in situ* caspase 3/7 activity, cell cycle progression was measured by BrdU incorporation. One sample t-test, data normalised to control for each assay. * p<0.05, **p<0.01, ***p<0.001, ****p<0.0001.

We sought to determine whether these observed reductions in growth and/or viability could be accounted for by induction of apoptosis. Proliferation was measured by resazurin reduction and BrdU incorporation, and apoptosis was assayed using an *in situ* fluorescent-on-cleavage Caspase 3/7 substrate, after 24 hours of drug treatment (**Figure 6C**). Doses of Samuraciclib, Dubermatinib and IPR-803 were chosen which gave comparable effects on attenuation of proliferation when assayed by resazurin reduction; proliferation was significantly decreased at all drug concentrations used. Apoptosis was not significantly induced at 0.25 and 0.5 µM doses of Samuraciclib, but was significantly increased with 1 µM Samuraciclib (p = 0.0003). The reduction in viability seen at lower doses of Samuraciclib, in particular the reduction in the proportion of cells entering S-phase, cannot be accounted for by apoptosis. Dubermatinib triggers a dose-dependent increase in apoptosis, while IPR-403 does not cause a large fold change in apoptosis at any concentration used.

## Discussion

Rapamycin can stabilise the accelerated loss of lung function suffered by LAM patients, but continued use is required, and some patients decline despite treatment [8]. Patients with the poorest lung function and longest disease duration at the start of treatment respond the least well to rapamycin [10]; histologically LAM lesions in these patients comprise both LAM cells and LAFs, and deposited ECM. Although in cancers with a dysregulated MTORC1 pathway rapamycin resistance is thought to be driven by the acquisition of additional mutations (reviewed in [32]) we hypothesise that in LAM the increasing complexity of the microenvironment of LAM cells provides paracrine signals which support LAM cell proliferation and survival in a rapamycin insensitive manner.

Our proteomic analysis of late-stage LAM lung tissue relative to healthy control tissue revealed upregulated components of the actin cytoskeleton, of glycolysis and of ECM. In the context of our model of LAM aetiology, these observations are consistent with accumulation of activated myofibroblasts (LAFs) in late-stage disease, since activated fibroblasts display enhanced glycolysis, intracellular stress fibres and produce increased amounts of extracellular matrix. Further, a recent study from our laboratory [33] in which a transcriptome of laser-captured LAM nodule tissue was generated, revealed significant upregulation of pathways including ECM organisation, focal adhesion, integrin signalling and ECM-receptor interaction.

We did not detect an increase in LAM cell markers, such as PMEL, VEGFD, or MMP11 [34] in LAM lung relative to healthy controls, which was unexpected, but could reflect a predominance of stromal fibroblasts at later stages of the disease as we have previously proposed [11]. Thus, although the LAM cell transcriptome from the LAM Cell Atlas [34] is consistent with a mesenchymal cell phenotype for LAM cells, including expression of *COL1A1*, *COL1A2*, *COL6A1, COL6A2*, and *COL6A3*, our data support the hypothesis that LAFs are the predominant source of deposited ECM in LAM lesions.

We reasoned that LAF-deposited ECM could be a paracrine factor driving LAM cell proliferation, and this proved to be the case in *in vitro* assays, using LAM patient-derived TSC2^-/-^ 621-101 cells. To better understand the response of LAM cells to ECM we used RNA sequencing to identify genes upregulated in 621-101 cells grown on LAF ECM relative to plastic, and mined these data for genes encoding known drivers of proliferation which were not downregulated by rapamycin treatment and which were potentially druggable - in common with other rare diseases, novel treatments for LAM are likely to come from repurposing existing drugs. We selected the upregulated genes *CDK7*, *GAS6* and *PLAU/PLAUR* for further investigation and chose to investigate whether inhibition of these pathways was able to attenuate the pro-proliferative effect of LAF ECM.

CDK7 is involved in both cell cycle regulation as a component of the Cyclin Activating Kinase (CAK) complex, and in transcription as a part of the core transcription factor TFIIH [35]. Transcription of *MNAT* (MAT1) and *CCNH* (Cyclin H), additional components of the CAK complex, was also elevated in cells grown on LAF ECM. Increased expression of CDK7 is found in several cancers [36]; and CDK7 inhibition has shown promise *in vitro* against prostate [37], pancreatic [38], gastric [39] and breast cancer cells [40]. Further, a covalent inhibitor of CDK7, THZ1, showed efficacy against *Tsc2*-null tumours in a mouse model [41]. We used the ATP-competitive CDK7 inhibitor Samuraciclib/CT7001; this drug is in clinical trials for advanced breast cancer [42], with a Fast Track designation from the U.S. Food and Drug Administration.

GAS6 is a high affinity ligand for the receptor tyrosine kinase AXL [43,44]. AXL is overexpressed in several cancers, is associated with poor prognosis, and is a promising target for anti-cancer therapies (reviewed in [45] and [46]). Further, AXL inhibition significantly reduced kidney tumour volume in a Tsc2^+/-^ mouse model [47]. To inhibit the GAS6/AXL axis we used both Dubermatinib/TP-0903 and LDC1267. Dubermatinib was developed clinically as an AXL inhibitor [48], but is better described as a multikinase inhibitor, with additional activity against ABL1, ALK, AURKA, AURKB, CDK4, CHEK2, FLT3, JAK2, JAK3, MERTK, PLK4, TYK2 and others, including CDK7 [49]. LDC1267 is not under clinical development but targets only members of the TAM family of kinases [30].

PLAU (urokinase) is a serine protease, which, when bound to its receptor, PLAUR (uPAR), cleaves plasminogen to plasmin and triggers a proteolytic cascade which activates ECM-degrading proteases and releases ECM-bound growth factors (reviewed in [50]). In addition, uPAR interacts with transmembrane receptors, including integrins [51], Epidermal Growth Factor Receptor [52] and Platelet-derived Growth Factor Receptor [53], activating downstream signalling pathways and potentially driving proliferation; these receptor-receptor interactions are potentiated by binding of urokinase to uPAR. Urokinase has previously been identified as upregulated in LAM lung tissue, and is critical for the progression of *Tsc2*^-/-^ tumours in a mouse model [54,55]. To inhibit urokinase and uPAR activity we used IPR-803, a small molecule inhibitor of the uPAR/urokinase interaction, in order to target both the protease activity of urokinase and uPAR-protein interactions potentiated by urokinase binding.

Inhibitors targeting each of these pathways were able to reduce LAM cell proliferation in *in vitro* assays in the presence of rapamycin, suggesting that these drugs may provide additional benefit to patients with LAM over rapamycin alone. However, we cannot be confident that the growth inhibitory activity of Dubermatinib against LAM cells was a consequence of AXL inhibition, since the more specific TAM kinase inhibitor LDC1267 failed to show an anti-proliferative effect in our experiments; further elucidation of the role of the AXL-GAS6 axis in LAM will require targeted genetic disruption of AXL. For Dubermatinib, the reduction in proliferation of treated cells could be accounted for by a dose dependent increase in apoptosis. For Samuraciclib at low doses, and IPR-803 at all doses used, apoptosis was not sufficient to account for the observed reduction in proliferation. Both drugs caused cell cycle exit or arrest, with Samuraciclib proving particularly potent in this respect.

Since CDK7 is a component of the transcription factor TFIIH, increased expression of CDK7 in response to ECM may mediate increased transcription of downstream pro-proliferative genes, making CDK7 a potential master regulator of ECM-driven proliferation and an attractive druggable target for LAM. Further, a recent report [29] demonstrated that active MTORC1 signalling, a hallmark of LAM cells, increases the sensitivity of cells to Samuraciclib, which triggers senescence. This is consistent with our observation that Samuraciclib triggers cell cycle arrest in TSC2^-/-^ model LAM cells.

In summary, we have confirmed our earlier observation that there is an accumulation of activated stromal fibroblasts in late stage LAM [11]. We initially proposed that these cells could support LAM cell proliferation and survival [5]; we have now identified one mechanism through which this support could be mediated, via the deposition of ECM by LAFs. Contact with LAF-deposited ECM elicits changes in gene expression in *TSC2*^-/-^ model LAM cells, including upregulation of expression of pro-proliferative genes. In this proof-of-concept screen we identified several promising pathways which could contribute to increased proliferation on LAF ECM, and showed that relevant pathway inhibitors attenuated proliferation to a greater extent than rapamycin alone. The pathways identified have all previously been linked with LAM or with the progression of *Tsc2*^-/-^ tumours in mouse models. Further investigation of both the nature of the interaction(s) between LAM cells and the ECM, and the mode of action of these inhibitors is required, however combining existing antifibrotic therapies, which target the deposition of ECM by fibroblasts, with drugs which target the response of LAM to ECM may provide new therapeutic options for patients with poor or partial response to rapamycin.

## Funding Statement

This work was supported by LAM Action, Registered Charity 1167610 (England & Wales), The LAM Foundation, the NIHR Nottingham Biomedical Research Centre and La Marató de TV3 Foundation.

## Acknowledgements

We are grateful to the women with LAM who consented to the use of their tissue samples and data in this work.

## Notes

### Competing Interest Statement

The authors have declared no competing interest.

