## Supplementary material for "Extracellular Matrix-Induced Genes May Reduce Response to Rapamycin in LAM": Figure Legends

### Supplementary Data Figure Legends

#### Figure 1

Histology of LAM lung tissue from five explanted lungs used for generation of shotgun proteome. LAM cells and LAFs are identified by alpha-Smooth Muscle Actin expression detected by immunohistochemistry. Scale bars = 2.5 mm

#### Table 1

Table showing gene names, p-values, and adjusted p-value of significant terms in the selected GO libraries interrogated with a list of 67 proteins that were more than 1.3 fold upregulated in LAM lung relative to healthy lung. The adjusted p-value is calculated using the Benjamini-Hochberg method for correction for multiple hypotheses testing. Only results with an adjusted p-value $<0.05$ are displayed.

#### Table 2

Genes differentially expressed between 621-101 cells grown on tissue culture plastic or LAF-deposited extracellular matrix (LAF ECM) for five days in medium containing 1% Foetal Calf Serum. Genes with a positive log2 fold change are upregulated by LAM ECM

#### Table 3

Genes differentially expressed between 621-101 cells grown on LAF-deposited extracellular matrix (LAF ECM) for five days in medium containing 1% Foetal Calf Serum in the absence (control) and presence of 10nM rapamycin. Genes with a positive log2 fold change are upregulated by rapamycin

#### Table 4

Genes differentially expressed between 621-101 cells grown on tissue culture plastic or LAF-deposited extracellular matrix (LAF ECM) for five days in medium containing 1% Foetal Calf Serum and 10 nM rapamycin. Genes with a positive log2 fold change are upregulated on LAF ECM with 10 nM rapamycin.

#### Table 5

Table showing gene names, p-values, and adjusted p-value of significant terms in the selected GO libraries interrogated with a list of 402 genes upregulated in 621-101 cells grown on LAF ECM relative to cells grown on plastic. The adjusted p-value is calculated using the Benjamini-Hochberg method for correction for multiple hypotheses testing. Only results with an adjusted p-value $<0.05$ are displayed.
