## Supplementary figures and images for "Extracellular Matrix-Induced Genes May Reduce Response to Rapamycin in LAM"

### Supplementary Figure 1

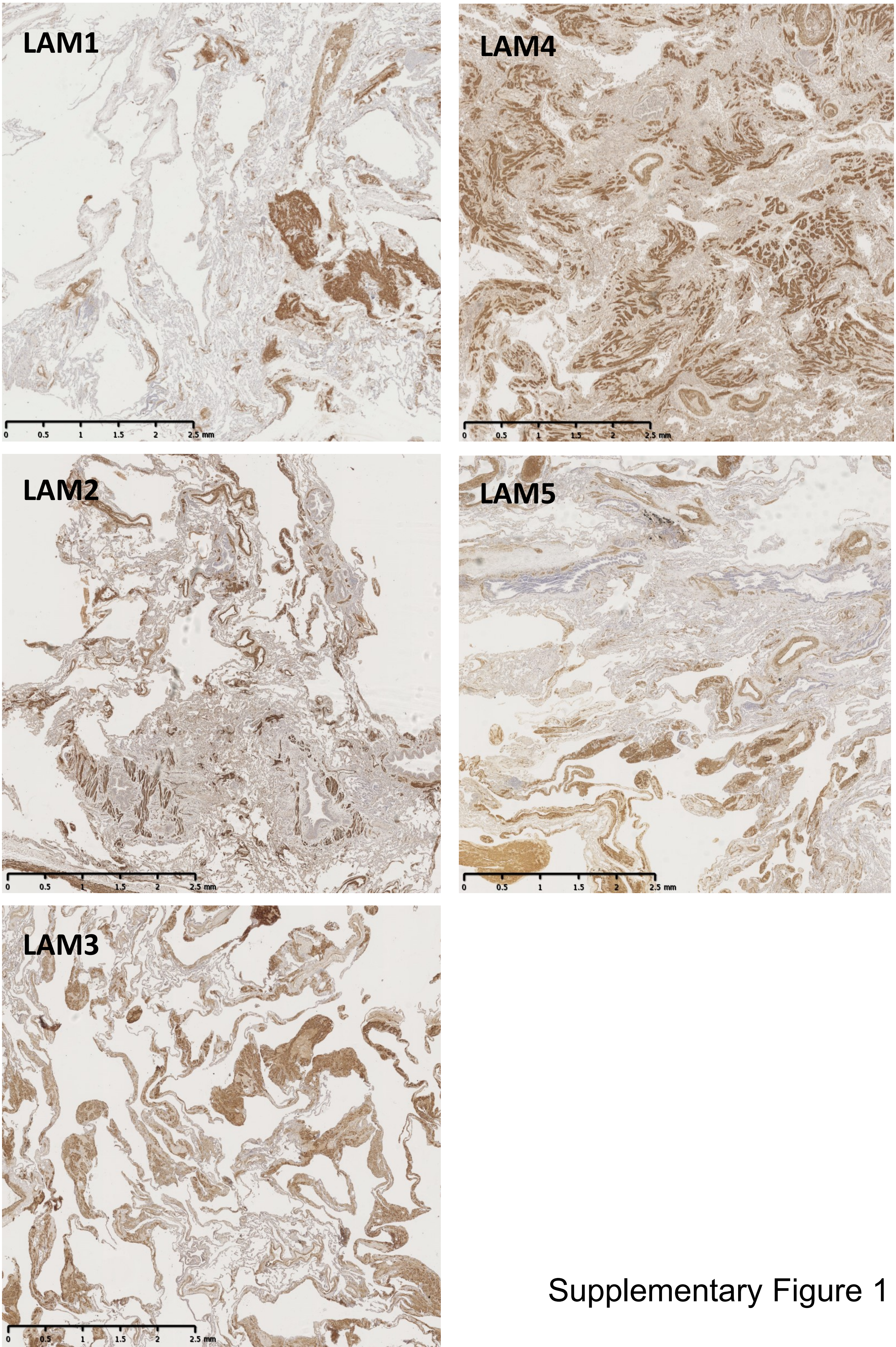


Supplementary Figure 1
